# *rahu* is a mutant allele of *Dnmt3c*, encoding a DNA methyltransferase homolog required for meiosis and transposon repression in the mouse male germline

**DOI:** 10.1101/121822

**Authors:** Devanshi Jain, Cem Meydan, Julian Lange, Corentin Claeys Bouuaert, Christopher E. Mason, Kathryn V. Anderson, Scott Keeney

**Author notes:** Corresponding authors, (DJ); (SK).

## Abstract

Transcriptional silencing by heritable cytosine-5 methylation is an ancient strategy to repress transposable elements. It was previously thought that mammals possess four DNA methyltransferase paralogs—*Dnmt1*, *Dnmt3a*, *Dnmt3b* and *Dnmt3l*—that establish and maintain cytosine-5 methylation. Here we identify a fifth paralog, *Dnmt3c*, that is essential for retrotransposon methylation and repression in the mouse male germline. From a phenotype-based forward genetics screen, we isolated a mutant mouse called ‘*rahu*’, which displays severe defects in double-strand-break repair and homologous chromosome synapsis during male meiosis, resulting in sterility. *rahu* is an allele of a transcription unit (*Gm14490*, renamed *Dnmt3c*) that was previously mis-annotated as a *Dnmt3-*family pseudogene. *Dnmt3c* encodes a cytosine methyltransferase homolog, and *Dnmt3c^rahu^* mutants harbor a non-synonymous mutation of a conserved residue within one of its cytosine methyltransferase motifs, similar to a mutation in human DNMT3B observed in patients with immunodeficiency, centromeric instability and facial anomalies syndrome. The *rahu* mutation lies at a potential dimerization interface and near the potential DNA binding interface, suggesting that it compromises protein-protein and/or protein-DNA interactions required for normal DNMT3C function *in vivo*. *Dnmt3c^rahu^* mutant males fail to establish normal methylation within LINE and LTR retrotransposon sequences in the germline and accumulate higher levels of transposon-derived transcripts and proteins, particularly from distinct L1 and ERVK retrotransposon families. Phylogenetic analysis indicates that *Dnmt3c* arose during rodent evolution by tandem duplication of *Dnmt3b*, after the divergence of the Dipodoidea and Muroidea superfamilies. These findings provide insight into the evolutionary dynamics and functional specialization of the transposon suppression machinery critical for mammalian sexual reproduction and epigenetic regulation.

## Author Summary

Half of human genomes are made up of transposons, which are mobile genetic elements that pose a constant threat to genome stability. As a defense strategy, transposons are methylated to prevent their expression and restrain their mobility. We have generated a mutant mouse, called ‘*rahu*’, that fails to methylate transposons in germ cells, suffers an increase in transposon expression and is sterile. *rahu* mice carry a mutation in a new gene, *Dnmt3c*, which appeared during rodent evolution through gene duplication 45–55 million years ago and is an essential component of the germline defense system against transposons in male mice.

## bioRxiv version 2 (Aug 7, 2017)

Two major experimental changes were incorporated in the revised manuscript:

1. We performed whole-genome bisulfite sequencing to map genome-wide the changes in methylation in *rahu* mutants. This analysis extended the characterization of the role of DNMT3C in transposon methylation.
2. To convince ourselves of the veracity of the *in vitro* DNA methylation results, we performed additional control experiments by purifying and assaying DNMT3C containing a mutation that is expected to completely eliminate methyltransferase activity (C537A). Unfortunately, this catalytically dead DNMT3C still gave a positive signal in the commercial methyltransferase activity assay we used. Thus, we are no longer able to say with certainty that we have been able to measure *bona fide* DNMT3C methyltransferase activity *in vitro*, and we are unable to draw firm conclusions about the effect of the *rahu* mutation on DNMT3C enzymatic activity. To be cautious, we have therefore removed the DNA methylation assay results from our revised manuscript and edited the text and figures accordingly.

## Introduction

Transposable elements have been described as ‘dark energy’ that acts both as a creative force, by giving rise to new genes and regulatory elements, and as a threat, by disrupting genome architecture [1]. Transposons occupy roughly 46% and 38% of the human and mouse genomes, respectively, and retrotransposons are the predominant class, including some that continue to be retrotransposition-competent [2-4]. Genomes have co-evolved with their retrotransposons and thus have multiple defense mechanisms to restrain transposon activity [4]. This restraint is of utmost importance in the germline, where retrotransposon activity not only facilitates vertical transmission, but also threatens genome integrity and germ cell viability.

The primary means of retrotransposon suppression is transcriptional silencing via cytosine-5 methylation [1, 5]. In the mouse male germline, after a developmentally programmed methyl-cytosine erasure, retrotransposon methylation is re-established in prospermatogonia prior to birth [6-9]. Shortly after birth, spermatogenesis initiates and spermatogonia enter a meiotic cell cycle, which encompasses an extended prophase. During meiotic prophase, the spermatocyte genome experiences developmentally programmed DNA double-strand breaks (DSBs) that are subsequently repaired by homologous recombination. Contemporaneously, homologous chromosomes align and build a proteinaceous structure called the synaptonemal complex (SC), which holds recombining chromosomes together (synapsis). Meiosis is completed by two successive divisions, forming haploid spermatids that enter the final developmental phase of spermiogenesis, culminating with the production of sperm [10]. Failure to methylate retrotransposons leads to abnormal retrotransposon expression, spermatogenic arrest in meiotic prophase, and sterility [11, 12].

Mammals possess three known enzymes that can catalyze DNA cytosine-5 methylation: DNMT1, DNMT3A, and DNMT3B. DNMT1 maintains DNA methylation by acting on hemimethylated DNA during replication, and *de novo* methylation is established by DNMT3A and DNMT3B [13-15]. The mammalian Dnmt3 family also includes DNMT3L, a catalytically inactive adaptor that lacks sequence motifs essential for cytosine-5 methylation [16-18]. DNMT3L interacts with and stimulates the activity of DNMT3A and DNMT3B [17, 19-23]. DNMT3L has also been implicated in the recognition of target sequences via interaction with histone H3 [24, 25].

Retrotransposon methylation in the germline is established by the concerted activities of DNMT3A, DNMT3B, and DNMT3L [7, 17, 26, 27]. Whereas mouse *Dnmt3a* and *Dnmt3b* are essential for development [15], *Dnmt3l*-deficient males are viable but sterile [17, 18]. They fail to methylate retrotransposons and accumulate retrotransposon-derived transcripts in both spermatogonia and spermatocytes. Abnormal retrotransposon expression is accompanied by defects in chromosome synapsis, inability to complete meiotic recombination, meiotic prophase arrest, and a complete absence of spermatids [11, 17, 27-29].

Here, we identify a new, fourth member of the Dnmt3 family in mice. Using a forward genetics approach, we isolated a male-sterile mutant that we named *rahu*, for *“*recombination-affected with hypogonadism from under-populated testes” (Rahu is a harbinger of misfortune and darkness in Vedic mythology). *rahu* mapped to a missense mutation in the predicted Dnmt3 pseudogene *Gm14490*. Re-named *Dnmt3c*, this gene is required for the methylation and repression of retrotransposons in the male germline. *Dnmt3c* was also independently discovered by Bourc’his and colleagues [30]; importantly, our results validate and expand their findings on the developmental role of *Dnmt3c*. We propose that *Dnmt3c* encodes a *de novo* DNA methyltransferase that diversified from DNMT3B during evolution of the Muroidea and that became functionally specialized for germline defense against retrotransposon activity.

## Results

### Isolation of the *rahu* meiotic mutants

We performed a random mutagenesis screen to recover mutants with autosomal recessive defects in meiosis. Male mice of the C57BL/6J (B6) strain were mutagenized with the alkylating agent *N*-ethyl-*N*-nitrosourea (ENU) [31-33], then a three-generation breeding scheme was carried out to “homozygose” recessive mutations, including outcrossing to FVB/NJ (FVB) mice to facilitate downstream genetic mapping (**Fig 1A**) [32, 34]. First, ENU-mutagenized B6 males were crossed to wild-type FVB females to generate founder males (F1) that are potential carriers of a mutation of interest. Then, each F1 male was crossed to wild-type FVB females to generate second-generation females (G2); if the F1 male was heterozygous for a mutation of interest, half of his daughters should be carriers. Finally, G2 daughters were crossed back to their F1 father to generate third-generation males (G3), of which one eighth should be homozygous for the mutation.

We screened juvenile (15–19 days *post-partum* (*dpp*)) G3 males for meiotic defects, focusing on recombination or its interdependent process, chromosome synapsis. To this end, we immunostained squashed spermatocyte nuclei for well-established meiotic markers whose deviations from wild-type patterns are diagnostic of specific defects: SYCP3, a component of axial elements and of lateral elements of the SC [10, 35, 36], and phosphorylated histone H2AX (γH2AX), which forms in response to meiotic DSBs [37]. As axial elements begin to form during the leptotene stage of meiotic prophase in wild type, SYCP3 staining appears as dots or short lines; axial elements elongate during zygonema and the first stretches of tripartite SC appear; SC juxtaposes autosomes all along their lengths in pachynema; and the SC begins to disassemble in diplonema (**Fig 1B**). Most DSBs are formed in leptonema and zygonema, resulting in florid staining for γH2AX at these stages, but this signal disappears from autosomes over the course of zygonema and pachynema as chromosomes synapse and DSBs are repaired by recombination (**Fig 1B**). γH2AX also accumulates in a recombination-independent manner in the sex body, a heterochromatic domain encompassing the sex chromosomes [37-40] (clearly visible in pachynema and diplonema in **Fig 1B**). Mutants with defects in recombination and/or SC formation deviate from these wild-type patterns in diagnostic ways [10, 36]. To streamline the screening process, we first used the SYCP3 staining pattern to classify meiotic prophase spermatocytes as either early prophase-like (leptonema or early zygonema) or later prophase-like (late zygonema, pachynema or diplonema). The γH2AX staining pattern was then evaluated to determine whether DSBs were formed and repaired properly and whether sex body formation appeared normal.

From F1 founder lines screened in this manner, we isolated a mutant line with SYCP3 and γH2AX patterns symptomatic of defects in meiotic DSB repair and/or synapsis (**Fig 1B and C**). Four of 28 G3 males from this line displayed a mutant phenotype in which spermatocytes with SYCP3 staining characteristic of early prophase I were abundant but later stages were absent or greatly depleted, indicating a block to meiotic progression (**Fig 1B and C and S1 Table**). Early prophase-like spermatocytes showed nucleus-wide γH2AX staining similar to wild type, consistent with formation of meiotic DSBs (**Fig 1B**). However, the mutants also had elevated numbers of abnormal spermatocytes displaying nucleus-wide γH2AX along with longer tracks of SYCP3 staining that were consistent with varying degrees of synapsis (**Fig 1B and C**). This pattern is a hallmark of recombination- and synapsis-defective mutants such as *Msh5^-/-^* and *Sycp1^-/-^* [36, 37, 41-43].

### rahu mutant males fail to complete meiosis, leading to male-specific infertility

Adult male *rahu* mutants were sterile, with none of the four animals we tested siring progeny. The mutant males displayed pronounced hypogonadism, with a 67% reduction in testes-to-body-weight ratio compared to littermates (mean ratios were 0.20% for *rahu* mutants and 0.62% for wild-type and heterozygous animals; p<0.01, one-sided Student’s t-test; **Fig 2A**).

**Figure 2.**
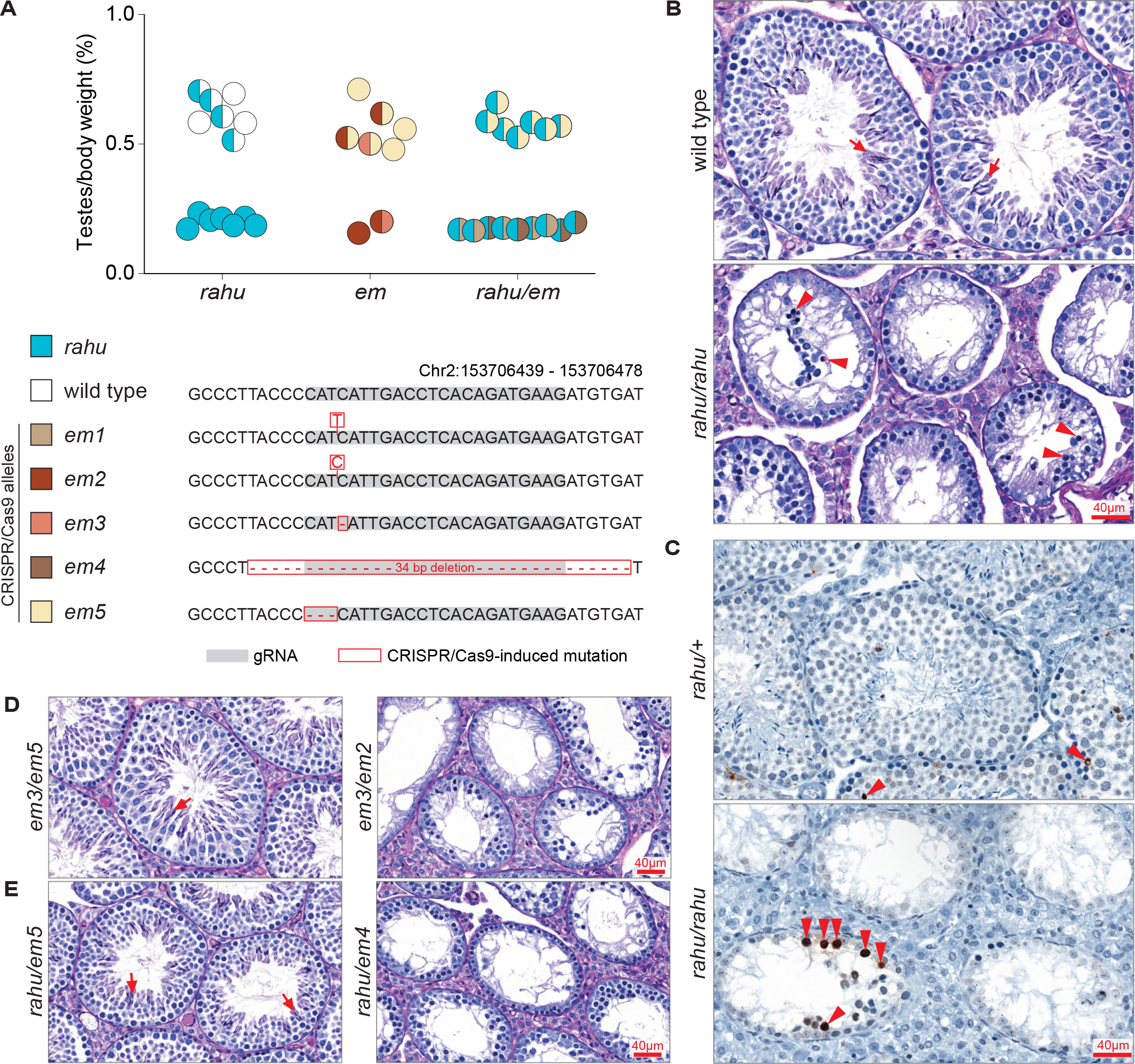
***rahu* and CRISPR/Cas9-targeted frameshift alleles of *Gm14490* cause meiotic arrest and fail to complement each other. (A)** The ratios of testes weight to body weight for 5- to 9-week-old mice carrying the *rahu* and CRISPR/Cas9-targeted alleles (*em*). Half-filled and fully filled circles represent heterozygous and homozygous genotypes, respectively. **(B)** Representative PAS-stained testis sections from adult mice of the indicated genotypes. Arrows indicate post-meiotic germ cells (spermatids) and arrowheads point to spermatocytes with an apoptotic morphology (condensed and/or fragmented). **(C)** Representative TUNEL-stained testis sections from adult mice of the indicated genotypes. Arrowheads point to TUNEL-positive cells (stained dark brown). **(D and E)** Representative PAS-stained testis sections from adult mice of the indicated genotypes. Arrows indicate post-meiotic germ cells.

In histological analysis of testis sections, seminiferous tubules from adult *rahu* mutants contained only early spermatogenic cells and completely lacked post-meiotic germ cells (**Fig 2B**). The most developmentally advanced cell types visible appeared apoptotic (**Fig 2B**), and increased apoptosis was confirmed by TUNEL staining (**Fig 2C**). The patterns suggest that spermatocyte arrest and apoptosis occur during mid-pachynema, possibly at stage IV of the seminiferous epithelial cycle, which is typical of mutants unable to properly synapse their chromosomes, repair meiotic DSBs, and/or transcriptionally silence their sex chromosomes [43-46]. Stage IV arrest is also observed in *Dnmt3l* mutants, which lack germline DNA methylation [17, 28].

In crosses with *rahu* heterozygous males, *rahu* homozygous females were fertile with an average litter size of 8.3 (19 litters from 10 dams), similar to *rahu* heterozygous females with an average litter size of 8.1 (29 litters from 10 dams). The *rahu* allele segregated in an expected Mendelian ratio: 26% wild type, 48% heterozygotes, and 26% mutant homozygotes from heterozygous dams (n=119); 49% heterozygotes and 51% mutant homozygotes from homozygous mutant dams (n=101). Homozygous mutants survived to adulthood and no morphological abnormalities have been apparent. We conclude that *rahu* does not severely impair female meiosis, embryonic development, or adult somatic functions.

### rahu is an allele of the novel gene Dnmt3c

We mapped the likely *rahu* causative mutation using a genetic polymorphism-based positional cloning strategy [34, 47]. Because the mutagenized B6 mice were outcrossed to FVB mice (**Fig 1A**), ENU-induced mutations should be linked to DNA sequence polymorphisms from the B6 background, so G3 males homozygous for recessive phenotype-causing mutations should also be homozygous for linked B6 polymorphisms. Furthermore, mutant mice from the same line should be homozygous for at least some of the same linked B6 polymorphisms, whereas related but phenotypically normal mice should not. We therefore searched for genomic regions of B6 single-nucleotide polymorphism (SNP) homozygosity that are shared between mutant *rahu* mice and not shared with phenotypically normal mice from the *rahu* line.

**Figure 1.**
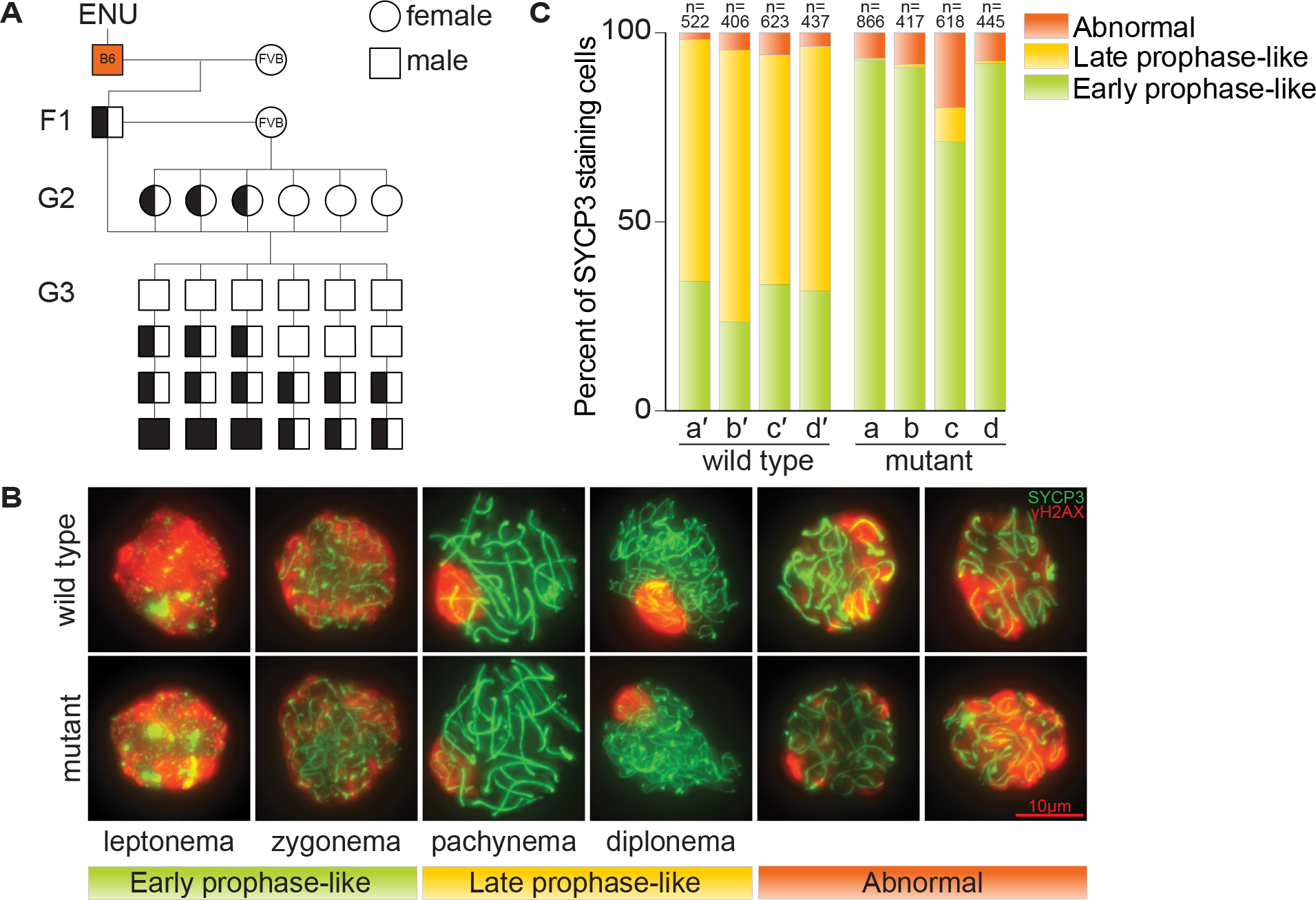
**Males from the ENU-induced mutant line *rahu* display meiotic defects. (A)** Breeding scheme used to isolate third-generation males with recessive defects in meiosis. Un-filled shapes represent animals that are wild-type for a mutation of interest, half-filled shapes are heterozygous carriers, and filled shapes are homozygotes. **(B)** Representative images showing the SYCP3 and γH2AX immunofluorescence patterns during meiotic prophase stages in squashed spermatocyte preparations from wild type and *rahu* mutants. Examples of cells with abnormal staining (nucleus-wide γH2AX along with longer tracks of SYCP3) are also shown. **(C)** Distribution of meiotic prophase stages in four G3 mutants obtained from the *rahu* line (a, b, c, d) and their phenotypically wild-type littermates (a′, b′, c′, d′). The number of SYCP3-positive spermatocytes counted from each animal is indicated.

We first coarsely mapped such regions by hybridization of genomic DNA from seven *rahu* mutants to a mouse SNP genotyping microarray. This yielded a single cluster of SNPs spanning 33.58 Mbp flanked by heterozygous SNPs *gnf02.126.027* (Chr2:127,800,747) and *rs3664408* (Chr2:161,380,222) (**Fig 3A**). Next, whole-exome sequencing of mutants identified seven un-annotated homozygous DNA sequence variants within the 33.58-Mbp mapped region (**Fig 3A**). To determine which was the likely causal mutation, we manually genotyped sequence variants within the 33.58-Mbp region in both mutant and phenotypically normal mice, targeting strain polymorphisms as well as the presumptive ENU-induced lesions themselves (**Fig 3A**). Presumptive ENU-induced lesions that were homozygous in phenotypically normal mice or that were heterozygous in meiosis-deficient mutants could be excluded as candidates. We applied this strategy to ~100 additional G3 and G4 mice, which allowed us to winnow the phenotype-causing mutation to within a 17.43-Mbp region flanked by the sequence change in the *Rrbp1* gene (Chr2:143,947,738) and SNP *rs3664408* (Chr2:161,380,222) (**Fig 3A**). This smaller region contained only one novel sequence variant, within a gene model named *Gm14490* (NCBI Gene ID: 668932 and Ensembl Gene ID: ENSMUSG00000082079).

**Figure 3.**
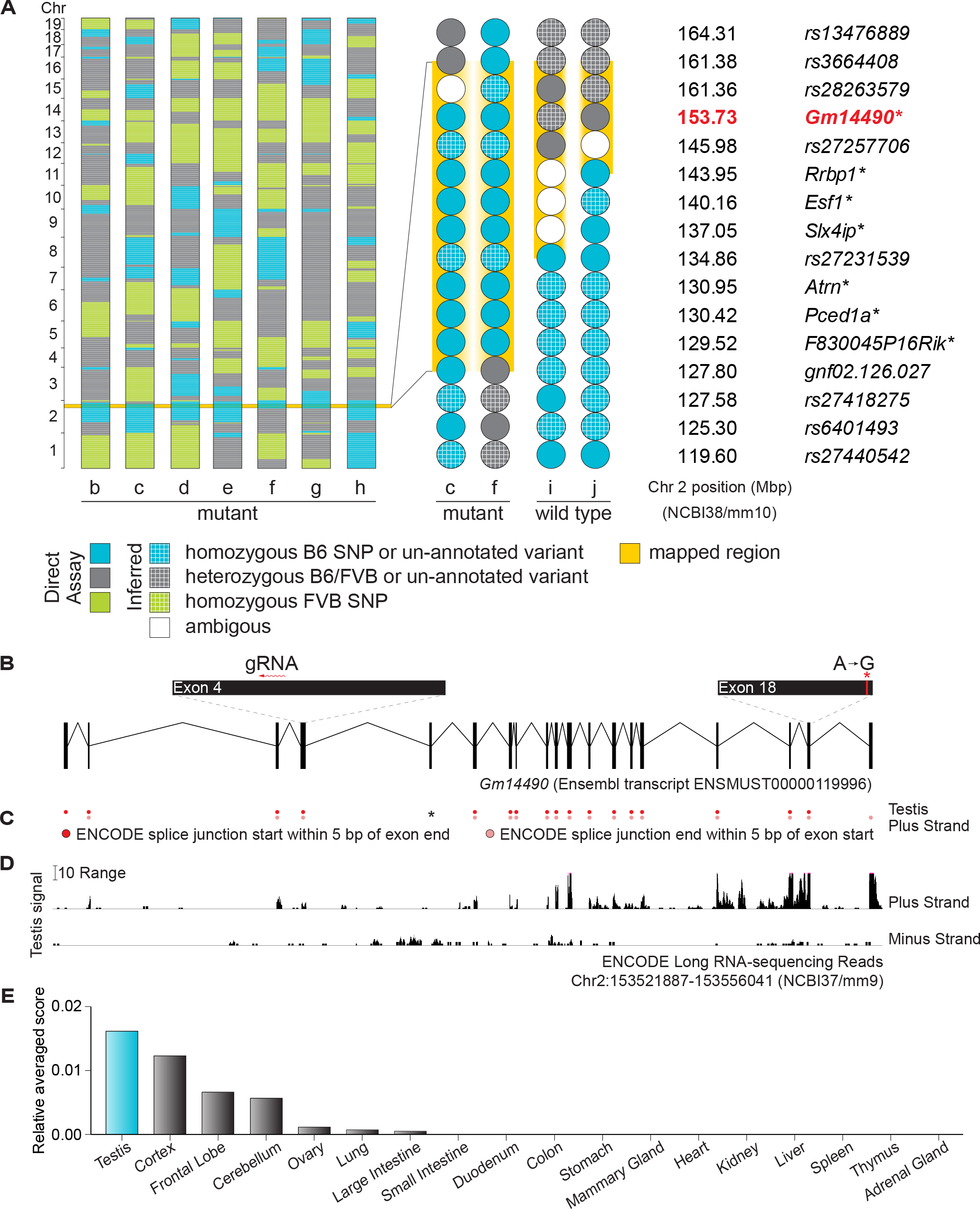
***rahu* is an allele of the testis-expressed gene *Gm14490*. (A)** SNP genotypes of seven *rahu* mutants (b, c, d, e, f, g, h) obtained using the Illumina Medium Density Linkage Panel are shown on the left. The single 33.58-Mbp region of B6 SNP homozygosity shared between mutants is highlighted in yellow. A detailed view of variants within this region is shown on the right for two informative *rahu* mutants (c, f) and two informative phenotypically wild-type mice (i, j). The reference SNP ID numbers (rs ID) for known variants and the gene names of previously un-annotated novel variants (i.e., presumptive ENU-induced lesions; asterisks) are listed. Phenotypes were assayed as shown in **Fig 1C** or **Fig 2A**. **(B)** Schematic of *Gm14490* (as predicted by Ensembl release 87) showing the locations of the ENU-induced lesion (red asterisk) and the gRNA used for CRISPR/Cas9-targeting. **(C)** Splice junctions that start or end within 5 bp of *Gm14490* exon boundaries, from ENCODE long RNA-sequencing (release 3) from adult testis. The black asterisk indicates a predicted exon with no splice junctions detected within 10 bp of its exon boundaries. **(D)** Density of mapped ENCODE long RNA-sequencing reads (release 3) from adult testis within a window spanning from 500 bp upstream of *Gm14490* to 500 bp downstream. The vertical viewing range of the displayed track is set at a minimum of 0 and maximum of 25; read densities exceeding this range are overlined in pink. **(E)** *Gm14490* expression estimate (ENCODE relative averaged score) in adult tissues.

The *rahu* lesion in *Gm14490* is an A to G nucleotide transition at position Chr2:153,727,342 (**Fig 3B**). Surprisingly, however, *Gm14490* was annotated as a pseudogene. It was first identified as a paralog of *Dnmt3b*, but the absence of expressed sequence tags and the presence of stop codons in the available gene build led previous researchers to conclude that *Gm14490* was a pseudogene [48]. However, more recent gene builds predicted a gene structure, encompassing introns with an open reading frame, that is matched by testis transcripts (**Fig 3B-D**). Mouse ENCODE RNA-sequencing data from adult testis revealed splice junctions within 5 bp of the boundaries for all exons except exon 5, suggesting that the predominant *Gm14490* transcript isoform(s) in adult testis does not contain exon 5 (**Fig 3B and C**). In the adult mouse, *Gm14490* is expressed in testis, with little or no expression in most somatic tissues other than brain (**Fig 3D and E**). *Gm14490* is predicted to yield a 2,218-nucleotide transcript (Ensembl Transcript ID: ENSMUST00000119996) containing 19 exons. The *rahu* point mutation is located in exon 18 (**Fig 3B**). The available expression data and the manifestation of a phenotype led us to surmise that *Gm14490* is not a pseudogene.

To test this hypothesis and to confirm that the identified point mutation is causative for the *rahu* phenotype, we generated targeted endonuclease-mediated (*em*) alleles of *Gm14490* by CRISPR/Cas9-mediated genome editing using a guide RNA (gRNA) targeted to exon 4 (**Fig 3B**). We analyzed four frameshift-generating alleles (*em1*, *em2*, *em3*, *em4*) and one in-frame deletion allele (*em5*), which results in a single-amino-acid deletion. We expected that frameshift mutations in an exon near the N-terminus of *Gm14490* are likely to lead to loss of function, whereas the single amino acid deletion would not. As predicted, young adult males carrying two copies of frameshift alleles had diminutive testes, similar to *rahu* mutants (**Fig 2A**). The in-frame deletion allele *em5* alone did not confer hypogonadism (homozygote mean = 0.55%, heterozygote mean = 0.57%). The *em2* homozygote showed reduced testes-to-body-weight ratios compared to compound heterozygotes also carrying the in-frame deletion *em5* allele (73% reduction; *em2* homozygote = 0.15%, *em2/em5* compound heterozygote mean = 0.57%), as did the *em2/em3* compound heterozygote (64% reduction; *em2/em3* = 0.20%, *em2/em5* and *em3/em5* compound heterozygotes mean = 0.55%).

Frameshift alleles did not complement the hypogonadism phenotype when crossed to *rahu* (**Fig 2A**). *rahu/em1* and *rahu/em4* compound heterozygotes had significantly reduced testes-to-body-weight ratios compared to *rahu/em5* compound heterozygotes (70% and 69% reduction, respectively; *rahu/em1* mean = 0.17%, *rahu/em4* mean = 0.17%, *rahu/em5* mean = 0.58%; p<0.01, one-sided Student’s t-test). Furthermore, adults with two frameshift alleles had depleted seminiferous tubules lacking post-meiotic germ cells, and the frameshift alleles did not complement the spermatocyte arrest phenotype when crossed to *rahu* (**Fig 2D and E**). Crosses between *em1/rahu* compound heterozygous females and *em1* heterozygous males gave litters (average size 9.3 pups; six litters from four dams). Also, *em2/em2* mutant females bred with *em2/+* heterozygote carrier males produced progeny (average litter size of six pups; two litters from two dams). Thus, similar to *rahu*, the frameshift mutations do not cause infertility in females.

We conclude that *rahu* is allelic to the *Gm14490* frameshift mutations and that *rahu* is likely to be a null or near-null for function of the gene. We further conclude that *Gm14490* is not a pseudogene and that it is essential during meiotic prophase in spermatocytes. On the basis of sequence homology to *Dnmt3b* and functional data shown below, we refer to *Gm14490* henceforth as *Dnmt3c*, also in keeping with a recent independent study [30].

### Dnmt3c encodes a likely DNA cytosine methyltransferase that is closely related to DNMT3B

*Dnmt3c* is predicted to encode a 739-aa protein with 77% overall similarity to the DNA cytosine methyltransferase DNMT3B (EMBOSS Water tool, Smith-Waterman algorithm [49-51]). It contains a cysteine-rich ATRX–DNMT3–DNMT3L (ADD) domain with 98.3% sequence similarity to that of DNMT3B, and a 96.8% similar DNA cytosine methyltransferase domain (**Fig 4A and S2 Table**). DNMT3C contains matches to both the sequence and arrangement of the six highly conserved cytosine methyltransferase domain motifs (I, IV, VI, VIII, IX and X) that are characteristic of active DNA methyltransferases (**Fig 4A and S2 Table**) [13, 52]. Among the mammalian Dnmt3 family members, DNMT3C shares most similarity with DNMT3B (**Fig 4B**), although it lacks a clear match to the Pro-Trp-Trp-Pro (PWWP) domain in DNMT3B (**Fig 4A**)[53]. The *rahu* mutation causes a glutamic acid to glycine substitution at position 692 within motif IX (**Fig 4A and C**). These data suggest that *Dnmt3c* encodes a novel DNA methyltransferase and that the *Dnmt3c^rahu^* mutant phenotype is a consequence of perturbing its methylation function in the male germline.

In a crystal structure of the cytosine methyltransferase domain of DNMT3A and the carboxy-terminal domain of DNMT3L, these peptides form a tetrameric complex with two DNMT3A and DNMT3L dimers, which further dimerize through DNMT3A-DNMT3A interaction [19]. Mutating residues at the DNMT3A-DNMT3A interface abolishes activity [54]. Homology-based modeling of the methyltransferase domain of DNMT3C places E692 near the potential dimerization interface (**Fig 4D and E**), as well as near the inferred DNA recognition region [19]. Thus, the E692G mutation in DNMT3C may interfere with protein-protein interactions and/or protein-DNA interactions that are required for normal DNMT3C activity.

**Figure 4.**
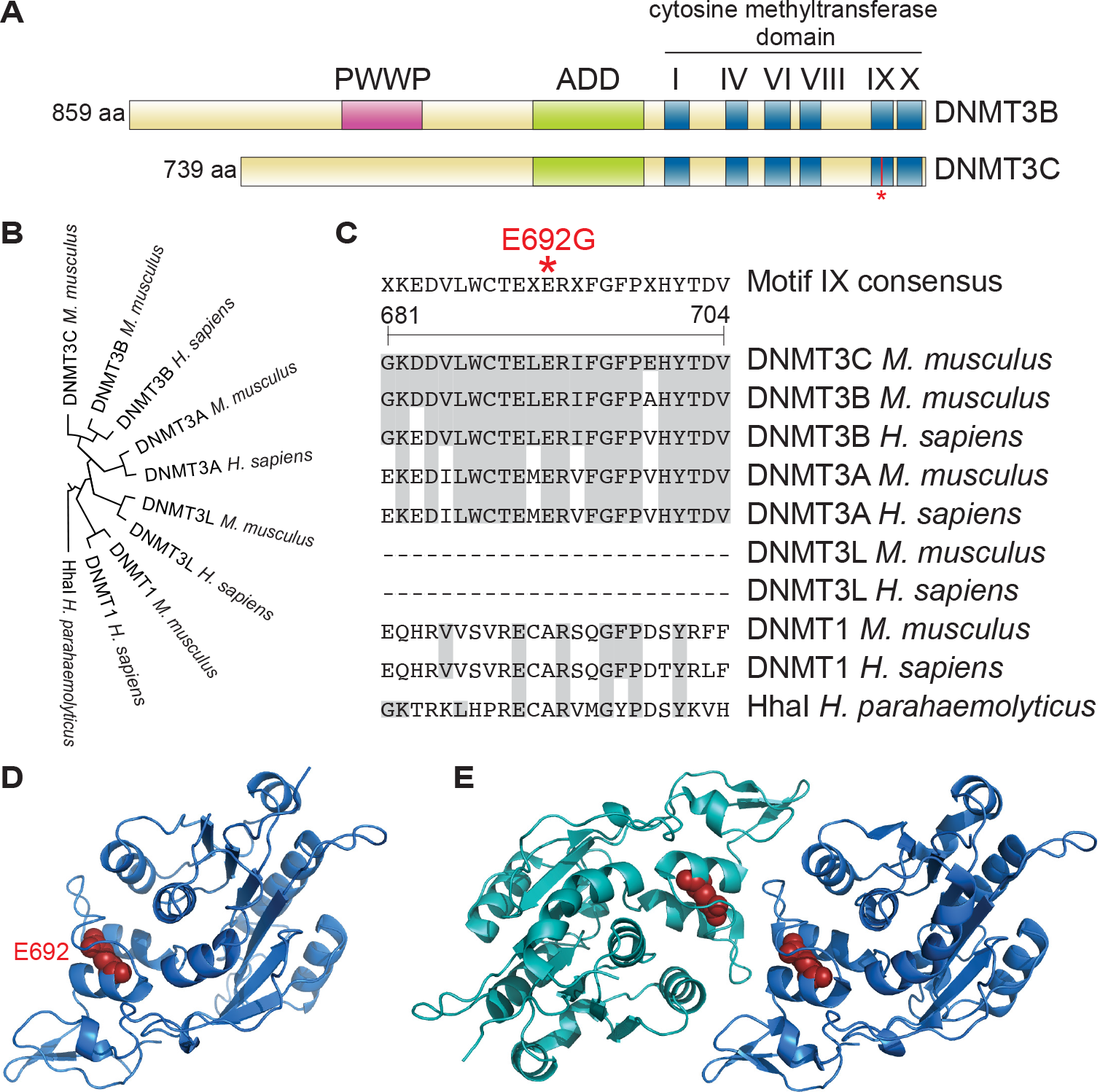
**DNMT3C is a putative DNA methyltransferase with similarity to DNMT3B. (A)** Schematics of DNMT3C and DNMT3B showing the location of conserved domains and the *rahu* mutation (asterisk). **(B)** Cladogram of Clustal Omega aligned human and mouse Dnmt3 family sequences rooted with HhaI. **(C)** Motif IX in Clustal Omega aligned sequences showing the location of the *rahu* mutation (asterisk). DNMT3L proteins do not contain Motif IX. Amino acids identical to those in DNMT3C are shaded gray. Amino acid positions refer to DNMT3C. **(D)** Homology-based model of DNMT3C cytosine methyltransferase domain with E692 depicted in red. **(E)** Crystal structure of the DNMT3A carboxy-terminal domain dimer (PDB ID:2QRV), with monomers depicted in two shades of blue. The DNMT3A amino acid equivalent to the glutamic acid that is mutated in *Dnmt3c^rahu^* mutants (DNMT3A E861) is shown in red.

### Dnmt3c is required for DNA methylation and transposon repression in the mouse male germline

A fundamental role of DNA methylation in the germline is to silence retrotransposons [27], so we examined the expression of long interspersed nuclear element-1 (L1) and intracisternal A particle (IAP) retrotransposon families, which are known to be active in the germline [5, 55-57]. Because the mutants undergo spermatogenic arrest, we examined animals at 14 dpp. At this age, testes predominantly contain spermatogonial stem cells and early meiotic cells, with the most advanced stage being pachynema [58]. *Dnmt3c^rahu^* mutants displayed increased expression of both L1 (~9-fold) and IAP (~3-fold) transcripts as assessed by quantitative RT-PCR (**Fig 5A**) (mean fold change of 8.5 using L1 ORFs primers and 8.7 using L1 ORF2 primers; mean fold change of 3.6 using IAP Gag primers and 2.1 using IAP 3′ LTR primers). We confirmed these findings by immunostaining testis sections for proteins encoded by L1 and IAP retrotransposons. In seminiferous tubules of heterozygotes, L1-encoded ORF1p was detectable at low levels and IAP-encoded Gag was barely detectable. However, both proteins accumulated to substantially higher levels in *Dnmt3c^rahu^* mutant testes (**Fig 5B**). Tubules of mutant juveniles contained L1 ORF1p in spermatocytes and IAP Gag in spermatogonia. Tubules from mutant adults contained prominent immunofluorescence signal for both proteins in spermatocytes. Thus, disruption of *Dnmt3c* derepresses expression of L1 and IAP retrotransposons in the male germline.

**Figure 5.**
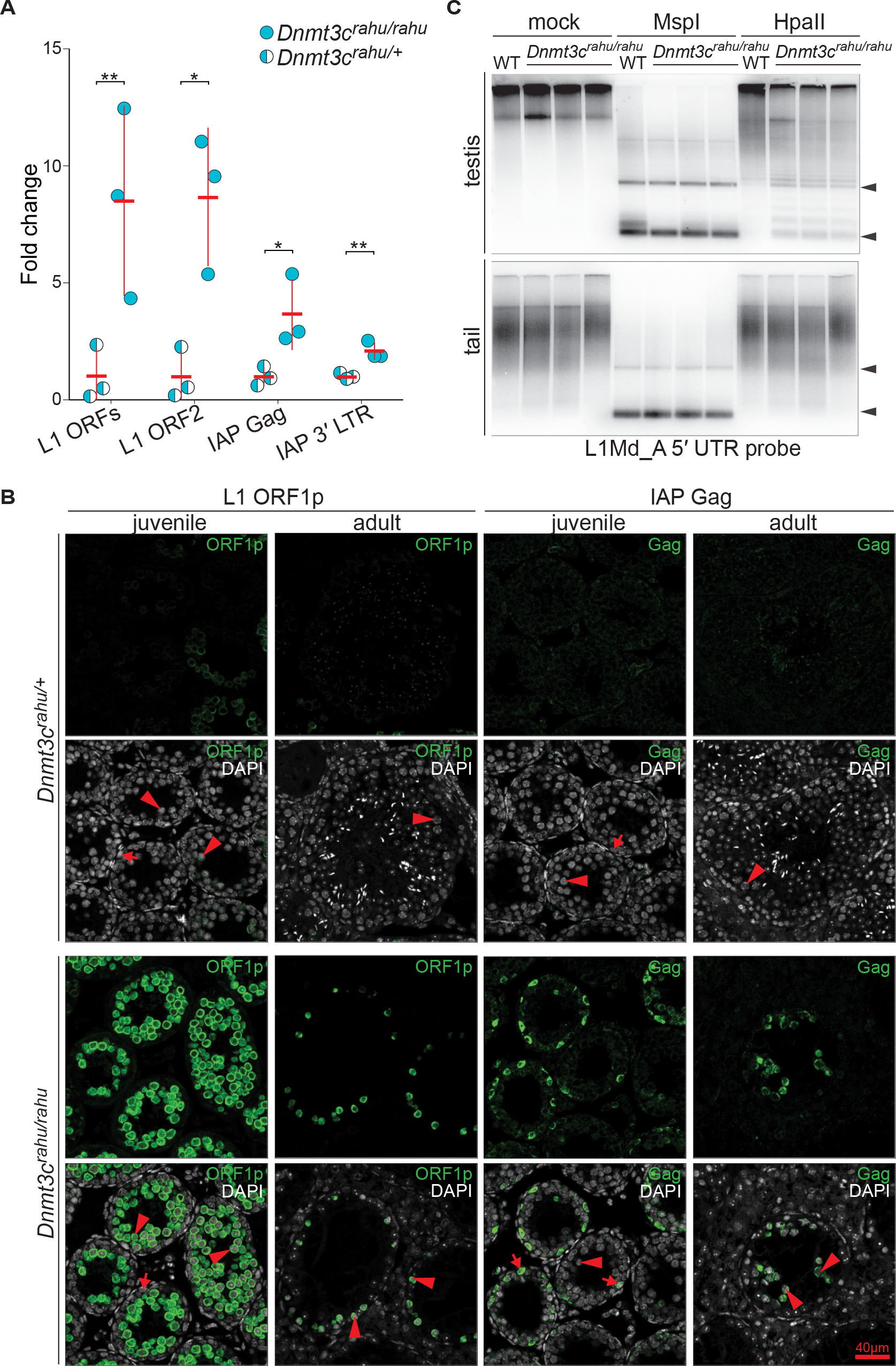
**Dnmt3c^*rahu*^ mutants exhibit phenotypes consistent with a role in DNA methylation and transposon repression in the male germline. (A)** Quantitative RT-PCR analysis of whole-testis RNA samples from six littermates (represented as individual data points) aged 14 dpp. Asterisk represents p<0.05 and double-asterisk represents p<0.01 in one-sided Student’s t-test. **(B)** Immunofluorescence of retrotransposon-encoded proteins L1 ORF1p and IAP Gag in testis sections from adults and from juveniles at 14 dpp. Matched exposures are shown comparing wild type with *Dnmt3c^rahu^* mutants. Arrows indicate spermatogonia and arrowheads point to spermatocytes. **(C)** Southern blot analysis of DNA extracted from either the testes or the tails of three *Dnmt3c^rahu^* mutants or a wild-type littermate at 15 dpp. DNA was digested with either the methylation-sensitive restriction enzyme HpaII, or its methylation-insensitive isoschizomer MspI. Arrowheads mark the positions expected for fully digested bands.

Next we assayed transposon DNA methylation directly by methylation-sensitive digestion and Southern blotting. Genomic DNA from juvenile *Dnmt3c^rahu^* mutants and wild-type littermates was digested with HpaII, for which DNA cleavage is blocked by CpG methylation within its recognition site, or with its methylation-insensitive isoschizomer MspI as a control. Digested DNA was separated on agarose gels and hybridized on Southern blots to a probe derived from the 5′ UTR of L1_MdA (A family of L1) sequences. DNA purified from somatic tissue (tail) was highly resistant to cleavage by HpaII, whether purified from homozyogous mutants or a heterozygous littermate (**Fig 5C**). This indicates that *Dnmt3c* is dispensable for DNA methylation in the soma, at least for L1Md_A elements. Testis DNA was also almost completely resistant to cleavage with HpaII when purified from heterozygotes, but substantially less so when purified from *Dnmt3c^rahu^* homozygous mutants (**Fig 5C**). We conclude that *Dnmt3c* is needed to establish normal levels of DNA methylation within L1Md_A elements, specifically in the germline. Taken together, these findings support a germline-specific function for *Dnmt3c* in retrotransposon DNA methylation and transcriptional repression.

### Dnmt3c represses transcription of distinct L1 and ERVK retrotransposon families

To more thoroughly assess the contribution of *Dnmt3c* in repressing retrotransposons, we performed RNA-seq on whole-testis samples from the same 14-*dpp*-old *Dnmt3c^rahu^* mutant and heterozygous littermates analyzed by RT-PCR. The expression levels of distinct LINE and LTR families were up-regulated in *Dnmt3c^rahu^* mutants, but SINE elements appeared to be changed little, if at all (**Fig 6A**). Because of variable expression between littermates of the same genotype, we analyzed the fold change of median expression values between mutants and heterozygotes (**Fig 6B**). Retrotransposons belonging to the L1 and ERVK superfamilies showed the strongest derepression. Specifically, L1 families L1Md_Gf, L1Md_T, L1_MdA and L1_Mm were up-regulated 12.1-, 9.6-, 6.4- and 2-fold, respectively. ERVK families IAPEZ-int, IAPLTR1_Mm, MMERVK10C-int, RLTR10C and IAPA_MM-int were up-regulated 10.1- to 4.5-fold. Two ERVK elements, IAPLTR3 and IAPLTR3-int were down-regulated in *Dnmt3c^rahu^* mutants (by 2.8- and 4.1-fold respectively).

The most affected L1 families in *Dnmt3c^rahu^* mutants are the same as those derepressed in *Dnmt3l* mutants, namely, young L1 families of the A, Tf and Gf subtypes [29, 59]. Comparison of the expression fold change in *Dnmt3l^−/−^* knockouts with *Dnmt3c^rahu^* mutants showed a striking overlap in effects for both the L1 and the LTR families (**Fig 6C**). This overlap is not an indirect effect of down-regulated *Dnmt3l* expression in *Dnmt3c^rahu^* mutants, as RNA-seq coverage for neither *Dnmt3l* nor *Dnmt3c* was significantly changed in *Dnmt3c^rahu^* mutants (*Dnmt3l* fold change 1.2, *Dnmt3c* fold change 1.0).

**Figure 6.**
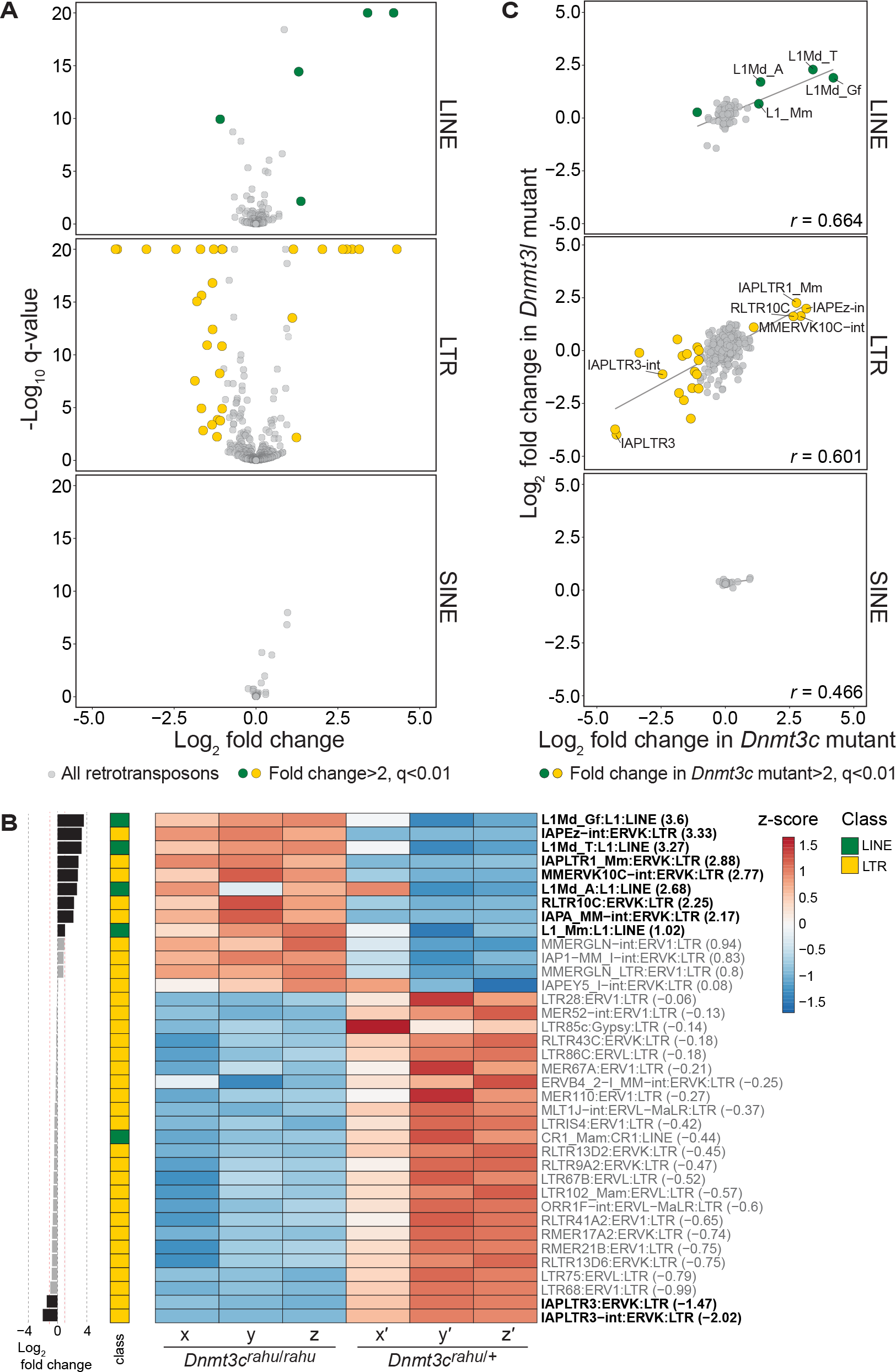
***Dnmt3c^rahu^* mutants up-regulate retrotransposons belonging to L1 and ERVK families. (A)** Volcano plot of differential RNA-seq values for various classes of retrotransposons. RNA-seq was performed on testis RNA samples from six 14-dpp animals from a single litter: three *Dnmt3c^rahu^* mutants and three heterozygotes (same mice analyzed in **Fig 5A**). Q-value is the Benjamini-Hochberg-adjusted p-value from DESeq2. Retrotransposons with expression fold change of >2 (up or down) and q < 0.01 are depicted as large, colored circles. **(B)** Heatmap showing the z-score of differentially expressed retrotransposon families (with expression fold change >2 and q < 0.01) in individual *Dnmt3c^rahu^* mutants (x, y, z) and their heterozygous littermates (x′, y′, z′). Labels on rows indicate the retrotransposon family, followed by superfamily, followed by class, and then in parentheses the log_2_ fold change of median expression in mutant versus heterozygote. Rows with greater than two-fold change in median expression (up or down) are in bold. The log_2_ fold changes are also provided in the bar graph at left (greater than two-fold change shown as black bars). **(C)** Correlation between differentially expressed retrotransposon families in 14-dpp *Dnmt3c^rahu^* mutants and 20-dpp *Dnmt3l* mutants. The regression line is shown and *r* is the Pearson correlation coefficient.

### Dnmt3c is required for methylation of L1, ERVK, and ERV1 retrotransposon families

To determine the global contribution of *Dnmt3c* to retrotransposons methylation, we performed whole-genome bisulfite sequencing (WGBS) on whole-testis samples from six 12-*dpp*-old *Dnmt3c^rahu^* mutant and wild-type littermates. CpGs within genes, CpG islands and SINE elements were mostly non-differentially methylated in *Dnmt3c^rahu^* mutants (**Fig 7A)**. An increase in the proportion of differentially methylated CpGs, specifically hypomethylated CpGs, was observed within LINE and LTR elements (16.04% and 6.62% hypomethylated CpGs, respectively) (**Fig 7A**). Meta-plots averaging over specific element types showed that LINE elements were on average hypomethylated near their 5′ ends, while LTR elements were hypomethylated on average across their entire bodies (**Fig 7B)**.

**Figure 7.**
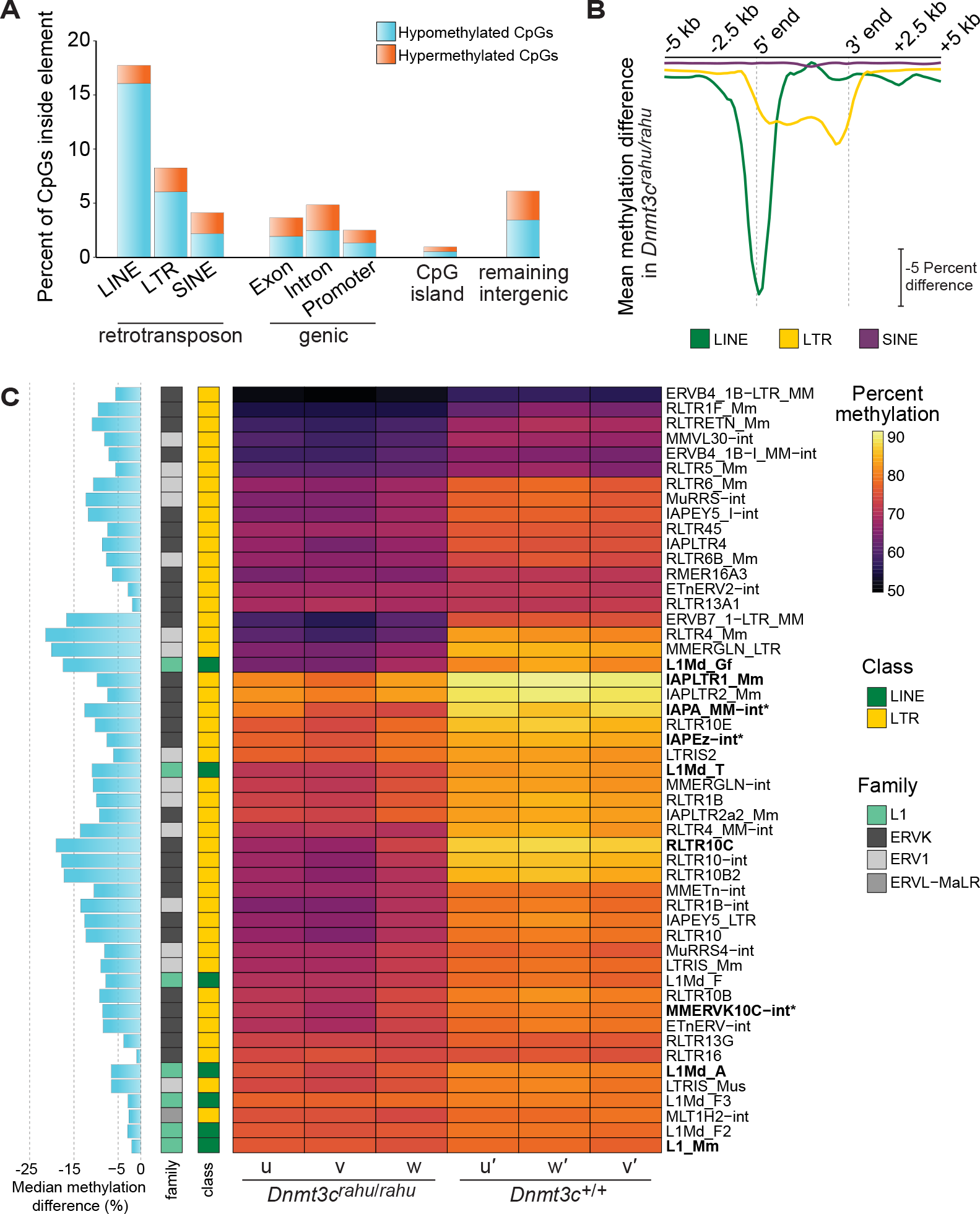
**Retrotransposons belonging to the L1, ERVK, and ERV1 families are hypomethylated in *Dnmt3c^rahu^* mutants. (A)** Proportions of differentially methylated CpGs within genomic elements, as determined by WGBS. WGBS was performed on whole-testis DNA samples from six 12-dpp animals (two *Dnmt3c^rahu^* mutant and two wild-type mice from one litter, and one *Dnmt3c^rahu^* mutant and one wild-type mice from a second litter). CpGs with >25% differential methylation (up or down) and with methylKit Sliding Linear Model-adjusted p-value <0.01 were considered differentially methylated. “Remaining intergenic” refers to genomic regions that do not overlap with LINE, LTR, SINE, genic, or CpG island annotations. **(B)** Meta-element plot showing the average difference in methylation levels between individual *Dnmt3c^rahu^* mutants and their wild-type littermates (mice u, v, u′, v′ from one litter; w, w′ from a second litter), across LINE, LTR, and SINE retrotransposons (including 5,000 bp of flanking sequence on each side) with minimum 95% coverage of the consensus sequence. **(C)** Heatmap showing the mean methylation levels of significantly changed retrotransposon families in individual *Dnmt3c^rahu^* mutants and their wild-type littermates (p < 0.01, two-sided Student’s t-test). Labels on rows indicate the retrotransposon family and differentially expressed retrotransposon families (families with greater than two-fold increase in median expression in *Dnmt3c^rahu^* mutants; rows in bold in **Fig 6B**) are in bold. Three differentially expressed retrotransposon families with changed methylation levels using a less stringent cutoff (p < 0.05, two-sided Student’s t-test) are included and are indicated with an asterisk. The median methylation difference is provided in the bar graph at left.

Comparison of the mean methylation levels of individual retrotransposon families showed that LINE elements belonging to the L1 superfamily and LTR elements belonging to the ERVK and ERV1 superfamilies were significantly hypomethylated in *Dnmt3c^rahu^* mutants (**Fig 7C**). In addition, one LTR element belonging to the ERVL-MaLR family, MLT1H2-int, was significantly changed. Consistent with a role for *Dnmt3c*-dependent DNA methylation in retrotransposon transcriptional repression, the specific retrotransposon families that were derepressed in *Dnmt3c^rahu^* mutants were also differentially methylated. For example, differentially expressed L1 families L1Md_Gf, L1Md_T, L1_MdA, and L1_Mm were significantly hypomethylated by 17.4%, 10.9%, 6.6%, and 2%, respectively (p <0.01, two-sided Student’s t-test) (**Fig 7C**). Differentially expressed ERVK families IAPLTR1_Mm and RLTR10C were significantly hypomethylated by 9.8% and 19.1%, respectively (p <0.01, two-sided Student’s t-test), and IAPEZ-int, MMERVK10C-int, and IAPA_MM-int were hypomethylated by 7.6%, 8.6% and 12.6%, respectively (p-value of 0.021, 0.025, and 0.020, respectively; two-sided Student’s t-test) (**Fig 7C, S1 Fig and S3 Table**).

### Dnmt3c arose by tandem duplication of Dnmt3b after divergence of the Dipodoidea and Muroidea rodent families

*Dnmt3c* is located directly adjacent to *Dnmt3b* in the mouse genome, and the two genes share 50.4% DNA sequence identity (EMBOSS Water tool, Smith-Waterman algorithm [49-51]). Dot-plot analysis of the genomic region encompassing both genes showed that the sequence identity shared between *Dnmt3c* and *Dnmt3b* extends over long stretches (up to 436 bp of 100% sequence identity and 2,805 bp of >95% sequence identity; appearing as horizontal lines in **Fig 8A**). Sequence similarity begins ~3,300 bp upstream of the annotated start of *Dnmt3c*, which matches the intronic region between *Dnmt3b* exons two and three. Similarity extends ~100 bp beyond the last annotated exon of *Dnmt3c* and matches the 3′ UTR of *Dnmt3b*, suggesting that these regions encode functional elements that have constrained sequence divergence (**Fig 8B**). *Dnmt3c* and *Dnmt3b* are remarkably similar with respect to their exon organization (**Fig 8B**), with stretches of sequence identity encompassing all *Dnmt3c* exons, except exons two and five. The sequence similarity in *Dnmt3c* is most extensive near its 3′ end (exons eight to nineteen), which encodes the ADD domain and methyltransferase motifs. As expected from the absence of the PWWP domain in DNMT3C, two of the three exons encoding this domain in *Dnmt3b* do not have obvious matches in the corresponding part of the genomic sequence of *Dnmt3c* (**Fig 8B**). Although *Dnmt3c* and *Dnmt3b* share extensive nucleotide identity, they are functionally specialized: whereas *Dnmt3b* establishes methylation both in somatic cells during embryogenesis and in the germline [7, 15, 26], *Dnmt3c* appears to function solely in the germline. Also, mouse and human DNMT3B protein sequences cluster separately from mouse DNMT3C (**Fig 4B**), suggesting that *Dnmt3c* and *Dnmt3b* have been evolving independently within the mouse lineage. These properties are consistent with a local gene duplication event followed by functional diversification.

**Figure 8.**
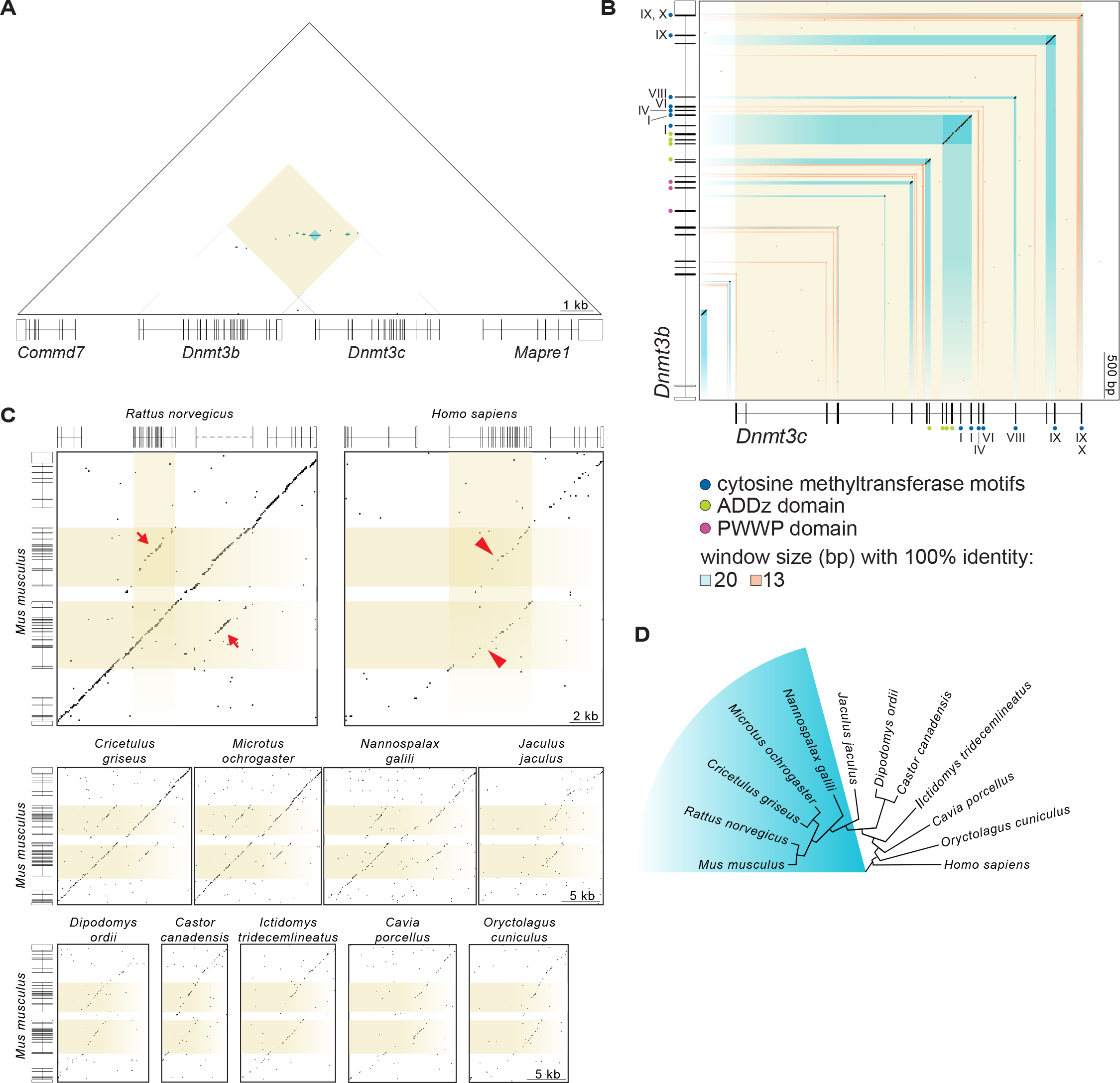
***Dnmt3c* arose by tandem duplication of *Dnmt3b* in rodents. (A)** Triangular dot plot of DNA sequence identities within a 156,377-bp region encompassing *Dnmt3b*, *Dnmt3c*, and flanking genes in *Mus musculus*. Each dt on the plot represents 100% identity within a 20-bp window. Direct repeats appear as horizontal lines. The yellow-tinted square shows the region within the plot that compares *Dnmt3b* to *Dnmt3c*, and the blue-tinted squares within it reflect regions with identical sequences within 20-bp windows. Immediately below the plot are gene models with shaded boxes representing coding sequences. **(B)** Dot-plot comparison of *M. musculus Dnmt3c* (including 3,500 bp of flanking sequence on each side) and *Dnmt3b*. Each black dot on the plot represents 100% identity within a 20-bp window, and blue-tinted rectangles highlight these regions. Each orange dot represents 100% identity within a 13-bp window, and orange-tinted rectangles highlight such regions when they are exonic or lie along the diagonal axis. Gene models annotated with exons encoding conserved domains are shown schematically along the axes. **(C)** Dot-plot comparisons of the *M. musculus* 156,377-bp region shown in (A) with its homologous region in other rodents (*Rattus norvegicus*, Norway rat; *Cricetulus griseus*, Chinese hamster; *Microtus ochrogaster*, prairie vole; *Nannospalax galili*, Upper Galilee Mountains blind mole rat; *Jaculus jaculus*, lesser Egyptian jerboa; *Dipodomys ordii*, Ord’s kangaroo rat; *Castor canadensis*, American beaver; *Ictidomys tridecemlineatus*, thirteen-lined ground squirrel; *Cavia porcellus*, domestic guinea pig), a lagomorph (*Oryctolagus cuniculus*, rabbit), and human (*Homo sapiens*). Each dot on the plot represents 100% identity within a 15-bp window. Yellow-tinted rectangles highlight *M. musculus Dnmt3b* and *Dnmt3c*, as well as *Dnmt3b* in rat and human. Gene models are shown for *M. musculus*, *R. norvegicus*, and *H. sapiens*. The putative *Dnmt3c* gene location in *R. norvegicus* is depicted by the gray dashed line above the dot plot. Segments of contiguous inter-species sequence identity between *Dnmt3b* and *Dnmt3c* appear as off-center partial diagonals (arrows) for those species that harbor the *Dnmt3b* and *Dnmt3c* pair, or as two offset diagonals (arrowheads) for species that lack the duplication. **(D)** Cladogram showing the evolutionary relationship of species analyzed (UCSC Genome Browser; [60]). Species that showed evidence of harboring *Dnmt3c* are highlighted in blue.

Dot-plot comparisons of the mouse genomic region containing *Dnmt3c* and *Dnmt3b* with homologous regions in rat and human showed that the duplication is also present in rat (appearing as a continuous central diagonal plus two off-center partial diagonals indicated by the red arrows in **Fig 8C**) but is absent in human (appearing as two offset diagonals marked by red arrowheads in **Fig 8C**). To determine the ancestry of this duplication event, we analyzed species from related rodent families (**Fig 8C and D**) [60] (UCSC Genome Browser). Dot-plot comparisons showed clear evidence of the *Dnmt3c* and *Dnmt3b* duplicate pair in Upper Galilee Mountains blind mole rat, Chinese hamster, and prairie vole, but not in lesser Egyptian jerboa or more distantly related rodents (**Fig 8C and D**). These findings indicate that the tandem duplication of *Dnmt3b* occurred between 55 and 45 million years ago, after the divergence of the Dipodoidea and Muroidea rodent superfamilies, but before the divergence of the Cricetidae and Muridae families [61].

## Discussion

This study illustrates the utility of forward genetic screens in identifying novel meiotic genes in mouse and reports the identification of *Dnmt3c*, a new gene essential for mouse spermatogenesis and genome regulation. We find that *Dnmt3c* functions in the methylation and subsequent repression of retrotransposons in the male germline, and that it arose by tandem duplication of the *Dnmt3b* gene. While this work was in progress, analogous findings were reported independently [30], and results from both studies agree well. Uniquely, we have isolated a novel point-mutated allele of *Dnmt3c* (*rahu*) that harbors a non-synonymous mutation of a conserved residue, and performed expression and epigenetic profiling of this mutant.

The presence of the six highly conserved cytosine methyltransferase motifs in DNMT3C and the methylation defect observed in *Dnmt3c^rahu^* mutants, which harbor a mutation within one motif (E692G), suggest that DNMT3C is enzymatically active. Indeed, DNMT3C methylates cytosines *in vitro*, and expression of *Dnmt3c* in *Dnmt1 Dnmt3a Dnmt3b* triple knockout embryonic stem cells leads to a gain in DNA methylation [30]. We found that homology-based modeling of the DNMT3C methyltransferase domain using the DNMT3A crystal structure places the E692 amino acid at a potential dimerization interface, as well as near the potential DNA binding interface [19]. Thus, the *rahu* mutation may compromise protein-protein interactions and/or protein-DNA interactions of DNMT3C. Intriguingly, a mutation affecting the corresponding region in DNMT3B is observed in patients suffering from immunodeficiency, centromeric instability and facial anomalies (ICF) syndrome, an autosomal recessive disease caused by mutations in DNMT3B [15, 62]. The patient-derived mutation results in a three-amino-acid (serine-threonine-proline) insertion immediately downstream of the corresponding glutamic acid that is mutated in *Dnmt3c^rahu^* mutants (DNMT3B Uniprot VAR_011502). Expression of recombinant mutant DNMT3B harboring this insertion in cell lines indicated that the mutation does not severely compromise protein stability. Rather, nuclear localization patterns were abnormal, suggesting that these residues may function in targeting DNMT3B to specific genomic regions [63]. We speculate that the E692G substitution in *Dnmt3c^rahu^* mutants may similarly interfere with DNMT3C localization to target sequences.

In mouse, *Dnmt3c* and *Dnmt3l* mutants have similar meiotic phenotypes, including male infertility due to spermatocyte arrest apparently occurring at epithelial stage IV ([17, 18, 27, 28, 30] and this study). Both genes are required for methylating and silencing retrotransposons in the male germline, and, consistent with previous findings, comparison of RNA-seq data from *Dnmt3c^rahu^* mutants with published *Dnmt3l^−/−^* data suggests considerable overlap in the transposon families that they target [27, 29, 30]. *Dnmt3l* is also required for methylation at imprinted loci [7, 17, 18, 26], but we did not observe embryonic defects characteristic of loss of methylation at maternally imprinted loci: *Dnmt3c* mutant females produced healthy litters of expected size, and their offspring survived to adulthood without discernible abnormalities, consistent with the findings of Barau *et al*. Among paternally imprinted loci, Barau *et al*. observed hypomethylation in *Dnmt3c* mutants only at the *Rasgfr1* imprinting control region. They also showed that *Dnmt3l* mutants were globally hypomethylated, including at intragenic and intergenic regions. In comparison, hypomethylation in *Dnmt3c* mutants is primarily restricted to L1, ERVK, and ERV1 retrotransposons ([30] and this study). Taken together, these results suggest that *Dnmt3c* provides a more specialized contribution than *Dnmt3l* to the germline methylation landscape.

The majority of differentially methylated regions in *Dnmt3c* mutants overlap with transposons ([30] and this study), so it is likely that the SC and γH2AX defects in *Dnmt3c^rahu^* mutants are indirect effects of transposon hypomethylation. Loss of transposon methylation in *Dnmt3l* mutants leads to abnormal levels of meiotic DSBs within transposon sequences, which in turn are thought to lead to deleterious non-allelic recombination events, culminating in meiotic arrest [29]. Similarly, transposon hypomethylation in *Dnmt3c^rahu^* mutants may result in an abnormal meiotic DSB landscape. A non-exclusive alternative is that the *Dnmt3c^rahu^* meiotic recombination defect may be linked to retrotransposon derepression, for example, via accumulation of DNA damage induced by a transposon-encoded endonuclease activity. This idea is supported by the presence of SPO11-independent DSBs in mice lacking *Maelstrom*, a piRNA pathway component required for transposon methylation and repression [64, 65]. *Maelstrom* mutant mice that also lack SPO11, whose catalytic activity is required for meiotic DSBs, show extensive immunofluorescence staining for DSB markers, consistent with damage that is mechanistically distinct from that induced during developmentally programmed meiotic recombination events. Yet another possibility is that transposon hypomethylation in *Dnmt3c^rahu^* mutants perturbs the expression of neighboring meiotic genes.

Comparative genomic analyses suggest that *Dnmt3c* arose in the Muroidea phylogenetic lineage by duplication of, and subsequent divergence from, *Dnmt3b* ([30] and this study). It is conceivable that *Dnmt3c* neo-functionalized in response to selective pressure imposed by an increase in the retrotransposon load within the genome. An alternative hypothesis is that in organisms that lack *Dnmt3c*, its function is performed by a *Dnmt3b* isoform or by a yet-to-be discovered Dnmt3 paralog. Given that the duplication is specific to muroid rodents and that *Dnmt3c* was previously mis-annotated as a pseudogene, its discovery exemplifies the power of forward genetic approaches. Moreover, the rapid evolution of meiotic proteins and the diversity of meiotic strategies adopted across different taxa necessitate organism-specific approaches. With advances in genomics facilitating the molecular characterization of phenotype-causing lesions identified in forward genetic screens, this approach will continue to be fruitful in furthering our understanding of gametogenesis.

## Materials and Methods

### Generation of Dnmt3c^rahu^ animals

All experiments conformed to regulatory standards and were approved by the Memorial Sloan Kettering Cancer Center (MSKCC) Institutional Animal Care and Use Committee. Male mice of the C57BL/6J background were mutagenized by three weekly injections of 100 µg ENU/g of body weight, then outcrossed to FVB/NJ females. Wild-type mice of both inbred strains were purchased from The Jackson Laboratory. The mutagenesis and three-generation breeding scheme to generate homozygous mutant offspring were conducted as described elsewhere [31, 34] (**Fig 1A**). To minimize the chance of repeated recovery of the same ENU-induced mutation, no more than ten F1 founder males were screened from each mutagenized male. Each F1 founder male was used to generate ≥six G2 females and ~24 G3 males.

For screening, testes from G3 males were dissected, snap-frozen in liquid nitrogen, and stored at −80°. Males were screened for meiotic defects at ≥15 dpp, by which age spermatocytes in the relevant stages of meiotic prophase I are abundant [58]. An upper age limit of 19 dpp was imposed to avoid the need for weaning. For a given line, spermatocyte squash preparation and immunostaining were carried out only after testes had been obtained from ~24 G3 males. This side-by-side analysis facilitated comparisons of phenotypes between mice. One testis per mouse was used to generate squash preparations of spermatocyte nuclei and immunostained with anti-SYCP3 and anti-γH2AX antibodies as described below. The second testis was reserved for later DNA extraction if needed. Based on the extent of axial element and SC formation, SYCP3-positive spermatocyte nuclei were classified as either early prophase-like (equivalent to leptonema or early zygonema in wild type) or late prophase-like (late zygonema, pachynema, or diplonema). The γH2AX staining pattern was then evaluated. For each immunostained squash preparation, we aimed to evaluate ~20 early prophase-like cells and ~50 late prophase-like cells, if present. Priority for subsequent mapping and further analysis was given to lines that yielded at least two G3 males with similar spermatogenesis-defective phenotypes, derived from two or more G2 females.

Genotyping of *Dnmt3c^rahu^* animals was done by PCR amplification using *rahu* F and *rahu* R primers (**S4 Table**), followed by digestion of the amplified product with Hpy188I (NEB). The *rahu* mutation (A to G) creates a novel *Hpy*188I restriction site.

### Generation of targeted Dnmt3c mutations

Endonuclease-mediated alleles were generated at the MSKCC Mouse Genetics Core Facility using CRISPR/Cas9. A guide RNA (target sequence 5′-CATCTGTGAGGTCAATGATG) was designed to target predicted exon 4 of *Gm14490* (NCBI Gene ID: 668932 and Ensembl Gene ID: ENSMUSG00000082079) and used for editing as described [66]. Using the T7 promoter in the pU6T7 plasmid, the gRNA was synthesized by *in vitro* transcription and polyadenylated, then 100 ng/µl of gRNA and 50 ng/µl of Cas9 mRNA were co-injected into the pronuclei of CBA × B6 F2 hybrid zygotes using conventional techniques [67]. Founder mice were tested for presence of mutated alleles by PCR amplification of exon 4 using *Gm14490* F1 and R1 primers (**S4 Table**), followed by T7 endonuclease I (NEB) digestion.

Mis-targeting of CRISPR/Cas9 to *Dnmt3b* was considered unlikely as the gRNA has 10 (50%) mismatches relative to the homologous region of *Dnmt3b* (in exon 7 of Ensembl transcript ENSMUST00000109774 (alignment of *Gm14490* exon 4 and *Dnmt3b* exon 7 using EMBOSS Water tool, Smith-Waterman algorithm [49-51]). Also, this *Dnmt3b* region lacks the protospacer adjacent motif (PAM). Nonetheless, we screened mice directly by PCR (*Dnmt3b* F1 and R1 primers; **S4 Table**) and T7 endonuclease I assay at the relevant region in *Dnmt3b* to rule out presence of induced mutations. Animals that were positive for *Gm14490* mutation and negative for *Dnmt3b* mutation were selected for further analysis.

We deduced the mutation spectrum of founder *Dnmt3c^em^* mice by PCR amplification of the targeted region from tail-tip DNA (*Gm14490* F1 and R2 primers; **S4 Table**) followed by Sanger sequencing (Seq1; **S4 Table**). Sequence traces were analyzed using TIDE [68], CRISP-ID [69], and Poly Peak Parser [70].

*Dnmt3c^em^* founder males mosaic for frame-shift mutations were bred to mutant *Dnmt3c^rahu^* females to generate compound heterozygotes carrying both the *Dnmt3c^rahu^* allele and a *Dnmt3c^em^* allele. *Dnmt3c^em^*-carrying founder mice were also interbred to generate homozygotes or compound heterozygotes carrying two distinct *Dnmt3c^em^* alleles.

Genotyping of *Dnmt3c^em^* animals was done by PCR amplification of the targeted region followed by Sanger sequencing. PCR amplification was done with either *Gm14490* F1 and R2 primers followed by sequencing with Seq1, or with *Gm14490* F2 and R2 primers followed by sequencing with Seq2 (**S4 Table**). Sequence traces were analyzed to determine the mutation spectrum as described above. *Dnmt3c^em^* animals were also genotyped for the homologous region in *Dnmt3b* by PCR amplification with *Dnmt3b* F2 and R1 primers followed by sequencing with Seq1 (**S4 Table**).

### Genetic mapping and exome sequencing

All genome coordinates are from mouse genome assembly GRCm38/mm10 unless indicated otherwise. For genetic mapping, the screen breeding scheme (**Fig 1A**) was expanded: additional G2 females were generated and crossed to their F1 sire, and were identified as mutation carriers if they birthed G3 males displaying the *Dnmt3c^rahu^* phenotype. Breeding of G2 carriers to the F1 founder was continued to accrue additional homozygous mutants.

The *Dnmt3c^rahu^* phenotype was coarsely mapped by microarray-based genome-wide SNP genotyping using the Illumina Mouse Medium Density Linkage Panel. To obtain genomic DNA, testes or tail biopsies were incubated in 200 µl of DirectPCR lysis reagent (Viagen) containing 1 µl of proteinase K (>600 mAU/ml, Qiagen) for 24 hr at 55°. DNA was subsequently RNase A-treated, phenol:chloroform-extracted, and ethanol-precipitated. Microarray analysis was performed at the Genetic Analysis Facility, The Centre for Applied Genomics, The Hospital for Sick Children, Toronto, ON, Canada. For bioinformatics analysis, 720 SNPs out of 1449 SNPs total on the linkage panel were selected based on the following criteria: autosomal location, allelic variation between B6 and FVB backgrounds, and heterozygosity in the F1 founder. For fine-mapping by manual genotyping of variants, genotyping was done by PCR amplification followed either by Sanger sequencing or by digestion with an appropriate restriction enzyme.

For whole-exome sequencing, DNA from three phenotypically mutant G3 mice was prepared as for microarray analysis and pooled into a single sample. Because the mutant mice should share the phenotype-causing mutation(s), we expected this pooling approach to boost the reliability of mutation detection. Whole-exome sequencing was performed at the MSKCC Integrated Genomics Operation. Exome capture was performed using SureSelectXT kit (Agilent Technologies) and SureSelect Mouse All Exon baits (Agilent Technologies). An average of 100 million 75-bp paired reads were generated. Read adapters were trimmed using FASTX-Toolkit version 0.0.13 (http://hannonlab.cshl.edu/fastx_toolkit/) and read pairs were recreated after trimming using a custom Python script. Reads were aligned to mouse genome assembly GRCm38/mm10 using Burrows Wheeler Aligner-MEM software version 0.7.5a [71] with default settings, and duplicate reads were removed using Picard tools version 1.104 (https://broadinstitute.github.io/picard/). A minimum mapping quality filter of 30 was applied using SAMtools version 0.1.19 [72]. Genome Analysis Toolkit version 2.8-1-g932cd3a (Broad Institute; [73-75]) was used to locally realign reads with RealignerTargetCreator and IndelRealigner, to recalibrate base quality scores using BaseRecalibrator, and to identify variants using UnifiedGenotyper with the following settings: mbq 17; dcov 500; stand_call_conf 30; stand_emit_conf 30. Variants were annotated using ANNOVAR software [76]. To obtain a list of potential phenotype-causing lesions, variants were filtered further to only include those that 1) had a minimum sequencing depth of six reads, 2) were called as homozygous, and 3) were not known archived SNPs (i.e., they lacked a reference SNP ID number). The positions of variants within the 33.5-Mbp mapped region that we identified using this strategy are as follows: Chr2:129,515,815 in *F830045P16Rik*; Chr2:130,422,084 in *Pced1a*; Chr2:130,946,117 in *Atrn*;Chr2:137,046,739 in *Slx4ip*; Chr2:140,158,720 in *Esf1*; Chr2:143,947,738 in *Rrbp1*; and Chr2:153,727,342 in *Gm14490*.

### ENCODE data analysis

ENCODE long-RNA sequencing data used are from release 3. We acknowledge the ENCODE Consortium [77] and the ENCODE production laboratory of Thomas Gingeras (Cold Spring Harbor Laboratory) for generating the datasets. GEO accession numbers are as follows: Testis GSM900193, Cortex GSM1000563, Frontal lobe GSM1000562, Cerebellum GSM1000567, Ovary GSM900183, Lung GSM900196, Large intestine GSM900189, Adrenal gland GSM900188, Colon GSM900198, Stomach GSM900185, Duodenum GSM900187, Small intestine GSM900186, Heart GSM900199, Kidney GSM900194, Liver GSM900195, Mammary gland GSM900184, Spleen GSM900197, Thymus GSM900192.

### Histology

For histology, testes isolated from adult or juvenile mice were immersed overnight in 4% paraformaldehyde (PFA) at 4° with gentle agitation, followed by two 5-min washes in water at room temperature. Fixed testes were stored in 70% ethanol for up to 5 days. Testes were embedded in paraffin, then 5-µm-thick sections were cut and mounted on slides.

The periodic acid Schiff (PAS) staining, immunohistochemical TUNEL assay, and immunofluorescent staining were performed by the MSKCC Molecular Cytology core facility. Slides were stained with PAS and counterstained with hematoxylin using the Autostainer XL (Leica Microsystems) automated stainer. The TUNEL assay was performed using the Discovery XT processor (Ventana Medical Systems). Slides were manually deparaffinized in xylene, re-hydrated in a series of alcohol dilutions (100%, 95% and 70%) and tap water, placed in Discovery XT, treated with proteinase K (20 µg/ml in 1× phosphate-buffered saline (PBS)) for 12 min, and then incubated with terminal deoxynucleotidyl transferase (Roche) and biotin-dUTP (Roche) labeling mix for 1 hr. The detection was performed with DAB detection kit (Ventana Medical Systems) according to manufacturer’s instructions. Slides were counterstained with hematoxylin and mounted with coverslips with Permount (Fisher Scientific). The immunofluorescent staining was performed using Discovery XT. Slides were deparaffinized with EZPrep buffer (Ventana Medical Systems) and antigen retrieval was performed with CC1 buffer (Ventana Medical Systems). Slides were blocked for 30 min with Background Buster solution (Innovex), followed by avidin-biotin blocking (Ventana Medical Systems) for 8 min. Slides were incubated with primary antibody for 5 hr, followed by 60 min incubation with biotinylated goat anti-rabbit (1:200, Vector Labs). The detection was performed with Streptavidin-HRP D (part of DABMap kit, Ventana Medical Systems), followed by incubation with Tyramide Alexa Fluor 488 (Invitrogen) prepared according to manufacturer’s instructions. After staining, slides were counterstained with 5 µg/ml 4′,6-diamidino-2-phenylindole (DAPI) (Sigma) for 10 min and mounted with coverslips with Mowiol.

PAS-stained and TUNEL slides were digitized using Pannoramic Flash 250 (3DHistech) with 20× lens. Images were produced and analyzed using the Pannoramic Viewer software. Immunofluorescence images were produced using a LSM 880 (Zeiss) with 40× lens.

### Cytology

Spermatocyte squashes were prepared as described [78], with few modifications. Isolated testes with the tunica albuginea removed were minced and suspended in 2% PFA in 1× PBS. Fixation was allowed for approximately 10 sec and spermatocytes were spotted onto slides. A coverslip was pressed down onto the spermatocytes to squash them, and the preparation was snap-frozen in liquid nitrogen. Slides were removed from liquid nitrogen and the coverslip was pried off, followed by three 5-min washes in 1× PBS with gentle agitation. Washed slides were rinsed in water, air dried and stored at −80°. For immunofluorescence, slides were thawed in 1× PBS for 5 min with gentle agitation and stained as described [79]. Slides were incubated with blocking buffer (1× PBS with 0.2% gelatin from cold-water fish skin, 0.05% TWEEN-20, 0.2% BSA) with gentle agitation for 30 min. Slides were stained with primary antibodies overnight at 4°, washed three times for 5 min in blocking buffer with gentle agitation, incubated with appropriate Alexa Fluor secondary antibodies (1:100; Invitrogen) for 30 min at room temperature, then washed three times for 5 min in blocking buffer. All antibodies were diluted in blocking buffer. Slides were rinsed in water and cover slips were mounted using mounting medium containing DAPI (Vectashield). Stained slides were stored at 4° for up to 5 days. Immunostained slides were imaged on a Marianas Workstation (Intelligent Imaging Innovations; Zeiss Axio Observer inverted epifluorescent microscope with a complementary metal-oxide semiconductor camera) using a 63× oil-immersion objective.

### Antibodies

Primary antibodies and dilutions used are as follows: mouse anti-SYCP3 (SCP-3 (D-1), 1:100, Santa Cruz, sc-74569), rabbit anti-γH2AX (p-Histone H2A.X (ser 139), 1:750, Santa Cruz, sc-101696), rabbit anti-ORF1p (1:1000, gift from A. Bortvin), rabbit anti-IAP Gag (1:5000, gift from B.R. Cullen).

### Domain annotation

DNMT3C protein sequence was obtained by translating the 2,218-bp Ensembl transcript ENSMUST00000119996 (release 87) cDNA sequence. An additional two nucleotides (AG) were added to the predicted transcript end to create a stop codon, resulting in a 2,220-bp transcript. DNMT3B protein sequence was obtained by translating the 4,320-bp Dnmt3b-001 Ensembl transcript ENSMUST00000109774.8 (release 87) coding sequence; this translation has 100% sequence identity to *M. musculus* DNMT3B with accession number O88509. ADD and PWWP domains were predicted using the NCBI Conserved Domain Database Search tool (accession numbers cd11728 and cd05835, respectively) [80]. To determine the cytosine methyltransferase motif locations in DNMT3C and DNMT3B, the following sequences were aligned by Clustal Omega alignment method using MegAlign Pro software (DNA STAR, Lasergene): *M. musculus* DNMT3C sequence determined as described above, *M. musculus* DNMT3B (accession number O88509), *H. sapiens* DNMT3B (accession number Q9UBC3), *M. musculus* DNMT3A (accession number O88508), *H. sapiens* DNMT3A (accession number Q9Y6K1), *M. musculus* DNMT3L (accession number Q9CWR8), *H. sapiens* DNMT3L (accession number Q9UJW3), *M. musculus* DNMT1 (accession number P13864), *H. sapiens* DNMT1 (accession number P26358), and *H. parahaemolyticus* HhaI (accession number P05102). Then the six highly conserved motifs (I, IV, VI, VIII, IX, X) were annotated as defined for HhaI [52]. The cytosine methyltransferase domain was annotated as the start of Motif I to the end of Motif X. The start and end locations for the domains and motifs are listed in **S2 Table**. Clustal Omega alignments from MegAlign Pro were used to produce a tree using BioNJ algorithm [81], and the figure was prepared using FigTree version 1.4.3 (http://tree.bio.ed.ac.uk/software/figtree). The model of DNMT3C was generated with Phyre2 protein structure prediction tool. The protein was modeled from template 2QRV (DNA methyltransferase domain of DNMT3A, 77% identity), with a confidence score of 100%.

### Methylation-sensitive Southern blotting

To extract genomic DNA, one half of a single testis or equivalent mg of tail tissue was incubated in 200 µl of DirectPCR lysis reagent (Viagen) containing 1 µl of proteinase K solution (>600 mAU/ml, Qiagen) for 24 hr at 55°. DNA was subsequently RNase A-treated, phenol:chloroform-extracted twice, and ethanol-precipitated. ~1 µg of DNA was digested for 4 hr at 37° with the methylation-sensitive HpaII restriction enzyme (NEB) or its methylation-insensitive isoschizomer MspI (NEB). ~250 ng of digested DNA was electrophoresed on a 0.9% agarose gel and transferred as described [82] to an Amersham Hybond-XL membrane (GE Healthcare). The L1 5′UTR probe has been described elsewhere [64] and corresponds to nucleotides 515–1628 of the L1 sequence (GenBank accession number M13002). The probe was random-priming labeled with [α^32^P]-dCTP using High Prime premixed solution (Roche). Hybridizations were carried out overnight at 65°.

### Quantitative RT-PCR and RNA sequencing

For RNA expression analysis, six littermates (three homozygous mutant and three heterozygous mice born from a cross between a *Dnmt3c^rahu/+^* male and a *Dnmt3c^rahu/rahu^* female) aged 14 dpp were analyzed. Procedures involving commercial kits were performed according to manufacturers’ instructions. Total RNA from one half of a single testis, after removing the tunica albuginea, was extracted using the RNeasy Plus Mini Kit containing a genomic DNA eliminator column (Qiagen). The RNase-Free DNase Set (Qiagen) was used to perform an on-column DNase digestion during RNA purification, as well as an in-solution DNase digestion after RNA purification, followed by RNA cleanup.

For RT-PCR, 1–3 µg of RNA was used with random hexamer primers to synthesize cDNA using the SuperScript III First-Strand Synthesis System (Invitrogen). cDNA was diluted five-fold or more for PCRs. Quantitative PCR was carried out using LightCycler 480 SYBR Green I Master (Roche) for detecting products on a LightCycler 480 II Real-Time PCR instrument (Roche). All reactions were done at 60° annealing temperature, with an extension time of 20 sec for L1 ORF2, IAP 3′ LTR and IAP Gag primers, and 40 sec for L1 ORFs primers (**S4 Table**). All reactions were done in triplicate and accompanied by control reactions using cDNA synthesized without reverse transcriptase (-RT controls). The success of reactions was confirmed by analysis of amplification curves, melting curves, and electrophoresis of representative amplification products on an agarose gel.

LightCycler 480 Software was used to quantify products by absolute quantification analysis using the second derivative maximum method. All crossing point (Cp) values were normalized to the mean Cp value obtained for triplicate *Actb* reactions, to get relative values. The mean of relative values for triplicate reactions was used to obtain mean relative values. The mean relative value represents the relative amount of product in any given mouse. To obtain fold change values, the mean relative value for each mouse was normalized to the mean of that obtained for three heterozygous mice.

RNA sequencing (RNA-seq) was performed at the Integrated Genomics Operation of MSKCC. 1 µg of total RNA underwent ribosomal depletion and library preparation using the TruSeq Stranded Total RNA LT kit (Illumina) with 6 cycles of post-ligation PCR. Sequencing was performed using Illumina HiSeq 2500 (2×100-bp paired-end reads) using the TruSeq SBS Kit v3 (Illumina). On average, 61 million paired reads were generated per sample. Resulting RNAseq fastq files were aligned using STAR version 2.4.0f1 [83] to the mouse genome (GRCm38/mm10) using Gencode M11 transcriptome assembly [84] for junction points. Coding genes and transposable elements were quantified using TEtoolkit [85] to annotate both uniquely and ambiguously mapped reads. Gencode annotation and Repbase database [86] for repetitive sequences and transposable elements were used during the quantification. Differentially expressed genes were calculated using DESeq2 [87] on the counts. For plotting, counts of transposable elements were normalized to all of the annotated reads including coding genes as counts per million (CPM). *Dnmt3l* RNA-seq data are published (GEO GSE57747 [29]). Expression values for 20-dpp-old *Dnmt3l* mutant and wild type, provided by the authors, were used to calculate retrotransposon expression fold change and matched to our RNAseq data by transposable element name.

### Whole-genome bisulfite sequencing

WGBS was performed with six animals from two litters (two homozygous mutants and two wild-type mice from one litter, and one homozygous mutant and one wild-type mouse from a second litter) aged 12 dpp. Genomic DNA was extracted from whole testis by incubating a single testis in 200 µl of DirectPCR lysis reagent (Viagen) containing 1 µl of proteinase K solution (>600 mAU/ml, Qiagen) for 24 hr at 55°. DNA was subsequently RNase A-treated, phenol:chloroform-extracted, and ethanol-precipitated.

Whole-genome bisulfite libraries were prepared and sequenced at the New York Genome Center (NYGC) using a tagmentation-based protocol developed by NYGC and J. Greally. In brief, 100 ng of genomic DNA was fragmented using tagmentation by Tn5 transposase and purified by Silane bead cleanup (Dynabeads MyOne Silane, Thermo Fisher Scientific). End filling was performed using dATP, dGTP, dTTP and methylated dCTP to protect added cytosines from bisulfite treatment. End-repaired DNA was purified by SPRI bead cleanup (Beckman), and subjected to bisulfite treatment and cleanup using EZ DNA Methylation-Gold MagPrep kit (Zymo Research). Illumina sequencing adapters (standard i5 adapter and custom i7 adapter) were added using PCR amplification. Finally, size selection of 300-bp to 800-bp fragments was performed using SPRI bead cleanup (Beckman). Libraries were sequenced on the Illumina X10 platform (v3 chemistry) using 2×150 cycles with standard Illumina read 1 primer and custom read 2 and i7 index primers to generate >375 million paired reads. Control library generated from *Kineococcus radiotolerans* was spiked at 20% to enhance library complexity.

Adapter sequence were first N-masked from raw FASTQ files using cutadapt v1.9.1 [88] (http://cutadapt.readthedocs.io/en/stable/index.html). Short-read alignment was performed with bwa-meth [89] (https://github.com/brentp/bwa-meth) to mouse genome assembly GRCm38/mm10. We modified bwa-meth’s default minimum longest match length for a read (0.44*read-length) to greater than 30 bp (0.2*read-length). The resulting alignment files were marked for duplicates using Picard v2.4.1 (http://broadinstitute.github.io/picard). Methylation levels were calculated using MethylDackel v0.1.13 (https://github.com/dpryan79/MethylDackel) at cytosines excluding quality control failed, supplemental, duplicate, and MAPQ less than 20 reads (which excluded multi-mapped reads). Additionally, bases with quality less than 20 or within the first 11 bases sequenced on either read pair were also excluded. For all downstream analyses, CpGs with minimum coverage of 5 reads were used. methylKit [90] was used to call differentially methylated CpGs with >25% differential methylation (up or down) and SLIM (Sliding Linear Model [91])-adjusted p <0.01. Differentially methylated CpGs were annotated using Repbase [86] for LINE, LTR, and SINE elements that have minimum 95% coverage of the consensus sequence. Refseq [92] was used to annotate genic regions and UCSC genome browser (http://genome.ucsc.edu) was used to annotate CpG islands. Methylation profile plots were made using elements that have minimum 95% coverage of the consensus sequence and contain minimum ten CpGs across the body of the element.

### Dot-plot and sequence analysis

Dot plots were made using custom Perl code available at http://pagelab.wi.mit.edu/material-request.html (David Page lab, Whitehead Institute, Cambridge, Massachusetts). A summary of sequences used is provided in **S5 Table**. All sequences were screened and masked for species-specific repetitive sequences prior to analysis using RepeatMasker software [93] with default settings. Two nucleotides (AG) were added to the annotated end of the *Gm14490* transcript to create a stop codon. Beaver, guinea pig, and rabbit dot plots were vertically reflected to maintain a consistent gene order of *Commd7* to *Mapre1* from left to right. Gene models were created based on Ensembl (version 87) exon annotations. To make the species cladogram, artificial sequences were aligned to resemble published data [60] (UCSC Genome Browser; http://genome.ucsc.edu) and the figure was prepared using FigTree version 1.4.3.

### Data Availability

Reagents are available upon request. RNA-seq data and WGBS data are available at Gene Expression Omnibus (GEO) with the accession number: GSE100960.

## Acknowledgments

We thank Alex Bortvin (Carnegie Institution of Washington) for providing the anti-L1 Orf1p antibody and the L1 probe; Bryan R. Cullen (Duke University Medical Center) for the anti-IAP Gag antibody; and Déborah Bourc’his for sharing information prior to publication. We thank Keeney lab members Luis Torres for assistance with variant mapping and mouse husbandry, and Diana Y. Eng and Jacquelyn Song for assistance with genotyping and mouse husbandry. We thank Nathalie J. Lailler (MSKCC Integrated Genomics Operation) for variant analyses of whole-exome sequencing data and Peter Romanienko (MSKCC Mouse Genetics Core Facility) for designing the gRNA. We thank the Genetic Analysis Facility (Centre for Applied Genomics, Hospital for Sick Children, Toronto, ON, Canada) for microarray analysis; the MSKCC Integrated Genomics Operation for whole-exome sequencing and RNA-sequencing; and the MSKCC Mouse Genetics Core Facility for CRISPR/Cas9-targeted mice.

---

**Figure S1.**
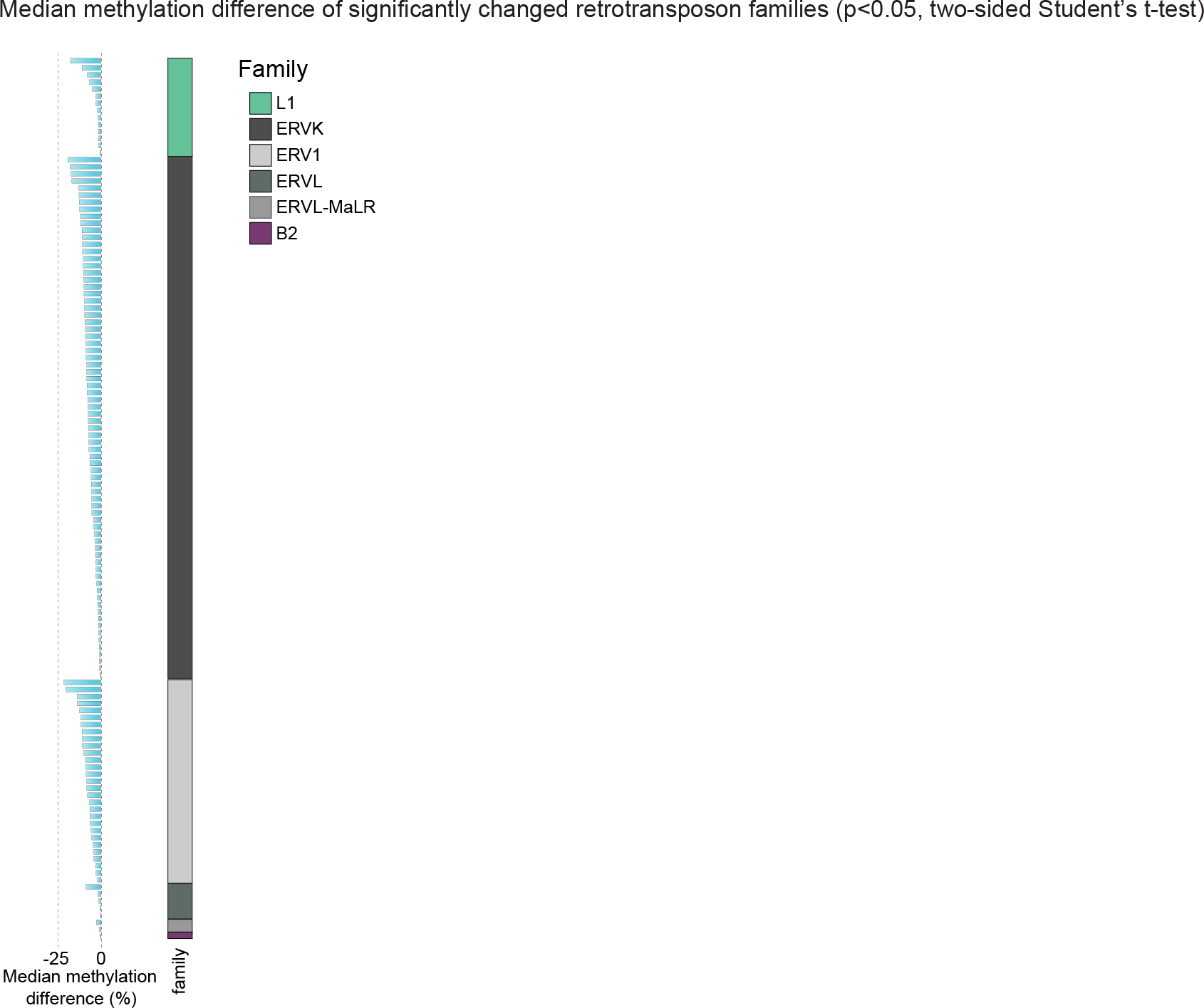
Median methylation difference of significantly changed retrotransposon families.

**Table S1.** *rahu* breeding results.

**Table S2.** DNMT3C and DNMT3B domain positions.

**Table S3.** Methylation levels of significantly changed retrotransposon families.

**Table S4.** Genotyping, cloning and quantitative RT-PCR primers.

**Table S5.** Dot-plot sequence coordinates.

